# Characteristics of vicarious touch reports in a general population

**DOI:** 10.1101/2024.01.30.577948

**Authors:** Sophie Smit, Matthew J. Crossley, Regine Zopf, Anina N. Rich

**Affiliations:** School of Psychological Sciences, Macquarie University, Sydney, Australia; Macquarie University Centre for Performance and Expertise, Macquarie University, Sydney, Australia; The MARCS institute for Brain, Behaviour & Development, Western Sydney University, Sydney, Australia

## Abstract

Vicarious touch broadly refers to sensory, neural, or affective responses to observing touch on another person. In this study, we focus on the conscious experience of these sensations. We collected vicarious touch reports from 422 undergraduate students who viewed videos from the Validated Touch-Video Database (VTD), a stimulus set systematically rated for hedonic qualities, arousal, and perceived threat levels. A surprisingly large majority (84%) of participants reported vicarious sensations such as touch, tingling, pressure, and pain, predominantly matching the location of touch on the hands observed in the videos. Contrary to previous studies, we found that vicarious touch was more prevalent in women than men. Additionally, while earlier research suggested a correlation between vicarious sensations and emotional reactivity, our data did not support this relationship. The nature and intensity of reported sensations varied strongly with the emotional content of the videos: pleasant touches often evoked ticklish or warm sensations, while unpleasant touches often resulted in pain or pressure. Participants regularly described their sensations as either pleasant or unpleasant, indicating that the affective qualities of observed touch were mirrored in their experiences. Subjective arousal and threat ratings taken from the VTD for each video strongly correlated with the prevalence and intensity of vicarious touch measured here, with painful, highly arousing touches eliciting the strongest responses. Even observing touch on a non-human object occasionally triggered similar vicarious sensations. Using a new clustering approach, we identified three distinct profiles of vicarious touch, each characterised by unique sensory experiences, variations in the localisation and intensity of sensations, and differences in responses to observing touch on another person versus an object. Overall, our results challenge the traditional view that conscious vicarious touch experiences are a rare phenomenon and underscore the diversity of vicarious sensory perception.

## Introduction

When watching a film where a character’s arm grazes a rough brick wall, some viewers may feel a corresponding sensation, as if their own arm had encountered the wall, while others might not react at all. The phenomenon of vicariously experiencing another person’s sensations, emotions, or actions is thought to be supported by a simulation mechanism, where observing—or inferring—another’s experience activates corresponding neural processes in the observer ^1–4^. While the term ‘vicarious touch’ is often used broadly to describe sensory, neural and affective responses to observed touch ^5,6^, here we specifically define it as the conscious experience of sensations when observing touch, acknowledging that this phenomenon includes both subthreshold activations and overt sensory experiences ^7,8^. Although ‘mirror-touch’ is sometimes used interchangeably with ‘vicarious touch’, it more commonly refers to synaesthesia-like experiences that involuntarily occur in daily life for a small proportion of people. Vicarious sensations are also reported in the general population, where experiences appear to be more variable and context-dependent ^9–14^, which is the focus of this study. Using the Validated Touch-Video Database (VTD) ^15^, we investigate how attributes of observed touch—such as subjective arousal, threat level, hedonic qualities, and the specific type of touch—shape vicarious sensations in a university sample.

Existing research has predominantly explored the subjective quality of vicarious touch through studies that prompt participants to report if they experienced any sensations in their own bodies after observing images or videos depicting touch involving another person. In such experimental contexts, ∼25-60% of participants report vicarious sensations such as touch, tingling, pressure, itchiness, or scratching ^4,9,10^. Furthermore, approximately one-third of participants report experiencing vicarious pain sensations, such as tingling, aching, or shooting pains, when witnessing others in painful situations, including injuries from injections or joint dislocations ^9–14^. While women report vicarious pain more frequently than men ^12,16^, no gender differences have been found in the reporting of vicarious non-painful touch ^9,10^. These findings suggest that vicarious sensations may be relatively common in experimental settings, though individual experiences can vary.

While existing research has suggested that reportable vicarious touch experiences may be quite common, it remains unclear what factors dictate the intensity and characteristics of these perceptions. For instance, research consistently shows that tingling is one of the most commonly reported vicarious sensations when observing touch or pain in others ^8,10,13^. If vicarious touch is the product of simulating the *actual sensation*, then tingling should be prevalent when observing pleasant, affective touch that would also produce tingling sensations if experienced directly ^17,18^. However, tingling is often reported even during observations of painful interactions ^13^, where it is presumably not a major element of the direct experience. Thus, it is currently unclear to what extent tingling can be ascribed to a sensory simulation process versus some other more generic process such as arousal or heightened bodily awareness ^19,20^.

Further exploration is also needed to fully comprehend how the emotional qualities of observed touch, such as perceived arousal, threat, and hedonic quality, influence subjective vicarious sensations. While functional Magnetic Resonance Imaging (fMRI) studies have investigated the various neural mechanisms involved in observing social, affective, pleasant, and unpleasant touches ^17,21–27^, it remains largely unclear if these varying observed touches also evoke differing *reportable sensations* in the observer. Previous self-report studies did find that observing a threatening touch—such as a knife tip touching a face—evoked stronger and more frequent sensations than benign touches like those from a fingertip or feather ^10,28^. Further investigation into how emotional stimulus characteristics shape subjective experiences can offer deeper insight into the nature and variability of vicarious touch.

Our understanding of vicarious touch has largely been shaped by studies on individuals with mirror-touch synaesthesia ^5,8–10,28–33^. This rare condition is characterised by frequent and intense localised tactile sensations, such as touch or pressure, rather than generalised sensations like tingling or itching ^8,10^. Individuals with mirror-touch synaesthesia also show a clearer distinction between observing touch on another person versus an object compared to non-synaesthetes ^10^, and reportedly exhibit enhanced social abilities, including heightened emotional empathy ^8,10,28,29^. However, distinguishing between mirror-touch synaesthesia and more common forms of vicarious touch remains challenging because they share overlapping sensory characteristics and comparatively little is known about vicarious touch experiences in non-synaesthetes – that is, people who do not report such experiences in daily life.

One intriguing possibility is that different vicarious touch profiles exist within the general population ^5^. For instance, some individuals may experience mostly localised tactile sensations, while others may perceive more generalised ones. These sensory differences may be accompanied by variations in sensation intensity, distinct responses to observed pain, differing reactions to observed touch on a person versus an object, and varying levels of empathy. Such tendencies could either distinctly categorise individuals, or vary along a continuum, with mirror-touch synaesthetes at the extreme end ^7,34–36^. Clustering approaches have been used to study vicarious pain profiles, showing that some individuals use predominantly localised and sensory descriptors (e.g., sharp, touch) for their experience, and others use more generalised and affective terms (e.g., ticklish, nauseating) ^12,16^. Using a clustering approach could similarly provide valuable insights into possible vicarious touch profiles.

In this study, we examine reported vicarious touch experiences in a large undergraduate sample using validated touch video stimuli from the VTD ^15^. These videos depict a close-up of a woman’s hands from a first-person perspective as she touches her own left hand with her right, either directly or using an object. We use these stimuli for several reasons. First, viewing touch from a first-rather than a third-person perspective may evoke stronger sensorimotor resonance, as it more closely aligns with one’s own bodily experiences ^37,38^. Second, observing a person touching themselves may engage different neural mechanisms than observing interpersonal touch ^39–41^, which can involve additional social-cognitive processing. These additional processes could alter vicarious sensations ^42^. Third, observed touch can be mapped using either an anatomical or specular reference frame ^28,39^. The VTD videos promote one type of mapping (i.e., anatomical, where touch on the left hand is more likely to be simulated on the observer’s own left hand), and minimises ambiguity introduced by third-person viewing, where touch may be mapped either anatomically (to the same body part) or specularly (to the opposite side in external space). By controlling for these perspective-related and social influences, our approach reduces variability and enables a more precise examination of how sensory and emotional stimulus characteristics influence vicarious touch responses.

Our work extends prior research by systematically investigating how key attributes of observed touch—including perceived arousal, threat level, hedonic qualities, and the specific nature of the touch—modulate vicarious sensations. In addition to assessing the presence of vicarious touch, we explore its qualitative characteristics, distinguishing between localised and generalised sensations and their relationship to stimulus properties. Our study is exploratory, addressing a broad range of questions about vicarious touch in the general population, using prior research to guide our analyses. For instance, findings suggest that women exhibit stronger responses to observed pain ^12,16,43,44^ but not to non-painful touch ^9,10^. To test for gender differences in vicarious sensations, we compared responses from men and women to observed touch with different hedonic qualities. Furthermore, the relationship between vicarious touch and general social abilities remains debated ^8,10,28,29,45^. Here, we investigate a possible link with emotional reactivity, a trait previously examined in the context of vicarious touch ^10,29,46^, which refers to an individual’s predisposition to share another person’s emotions. We also employ a data-driven clustering approach to identify subgroups of vicarious responders, offering novel insights into individual differences. In sum, we explore how sensory and emotional-affective stimulus characteristics influence vicarious touch experiences and how these relate to individual differences in the general population.

## Materials and Methods

### Participants

We recruited naïve undergraduate students via the Macquarie University subject pools and participants received course credit for their participation (N = 422; 292 women, 129 men, 1 non-binary; age *M* = 21.17 years, *SD* = 5.77 years, range = 17-56 years). As we only had one non-binary participant, we could not include their data in our analyses of gender. Students who had previously participated in our study for validating touch videos from the VTD ^15^ were not eligible to participate, as familiarity with the visual stimuli could potentially influence their experience. Written informed consent was obtained from all the participants. The study was approved by the Macquarie University Human Research Ethics committee (reference number: 52020925922588) and all methods were performed in accordance with the relevant guidelines and regulations of the Macquarie University ethics committee and the Declaration of Helsinki.

### Stimuli

We used stimuli from the VTD ^15^, consisting of 90 videos depicting self-touch interactions, where a woman touches her own left hand with her right hand, either directly or using an object. The hands are shown against a dark backdrop from a first-person perspective, with the left palm facing down. These videos, previously validated by an independent group of participants, display varying levels of arousal, threat, hedonic qualities, and different types of touch. Using the validation data, we selected ten videos representing each hedonic category (neutral, pleasant, unpleasant, painful; 40 total) for the current study. Further, we included five additional (non-validated) videos showing tactile interactions with an object (white block). These object videos were filmed under the same conditions as the hand videos and depict stroking the object with a soft brush, touching the object with a knife, tapping the object with a finger, stroking with a plastic brush next to the object, and tapping with a finger next to the object.

### Experimental procedure

After participants signed up to our study, they were given a link to an online questionnaire hosted on Qualtrics. We instructed participants to complete the task remotely on their own devices and to use a desktop computer or laptop instead of a phone or tablet. The questionnaire included the 45 touch videos presented in randomised order. To ensure that participants were ready and paid full attention to the entire video, we disabled auto-play, and the first frame of each video was a white screen with the text “Press play to start video”. After watching each video, participants were asked the following questions:

Q1: *While watching the video, did you experience a tactile sensation on YOUR body?* [Yes/No]. If participants answered ‘No’, they would go on to the next video. If they answered ‘Yes’, they would be presented with these follow-up questions:
Q2: *Which of these options describes the sensation on YOUR body?* (multiple answers allowed) [answer: Touch, Pain, Tingling, Pressure, Scratching, Ticklish, Warm, Cold, Other (describe)]
Q3: *How intense was it?* [answer: 1 = not at all, 10 = extremely]
Q4: *How did it feel?* [single answer out of the following options: neutral, pleasant, unpleasant, painful]
Q5: *How (pleasant/unpleasant/painful; only one category was presented and depended on the previous answer) was the touch?* [answer: 1 = not at all, 10 = extremely]
Q6: *Where on YOUR body did you feel the tactile sensation?* (multiple answers allowed) [answer: Left hand, Right hand, Another body part (describe), A general feeling (describe), Other (describe)]

Q2 allowed participants to select multiple sensory descriptors to characterise any vicarious sensations. Predefined options, based on prior studies on vicarious touch and pain, covered commonly reported experiences ^10–12,16^, while an ‘other’ option ensured participants were not restricted to these choices. Q4 assessed the hedonic quality of touch, which shapes affective responses at both neural and behavioural levels ^47,48^. In our videos, this ranged from pleasant (e.g., gentle caresses) to aversive (e.g., pain). While both pain and unpleasant touch are aversive, pain is typically associated with sharp or intense sensations, whereas rough textures or sustained pressure may feel unpleasant without being painful ^49,50^. Pain was included in both Q2 and Q4 to capture distinct aspects of participants’ responses: Q2 provided a broad characterisation of sensory qualities, while Q4 required a single categorical rating (neutral, pleasant, unpleasant, painful) to assess overall affective impact.

After finishing this part of the survey, participants completed five questions from the emotional reactivity subscale of the short Empathy Quotient (EQ) ^51^. Emotional reactivity has been linked to vicarious touch experiences in prior research ^10,29,46^, making this subscale an appropriate choice for our study. This validated short version ^51^ of the original EQ ^52^ was selected to minimise participant burden. The entire questionnaire took between 10 and 30 minutes to complete depending on how often the participant responded ‘Yes’ to the first question (as this resulted in five additional questions for each video).

### Analyses

Our analyses can be divided into three sections. In the first section, we investigate the general characteristics of vicarious touch in our population sample, exploring the prevalence of reported vicarious sensations, any gender differences, and the potential link with emotional reactivity. In the second section, we look at the effects of observed affective qualities (arousal, threat, hedonic quality) and the type of touch in the videos (e.g., stroking versus pressing) on the prevalence, intensity, and phenomenology of various touch experiences. In the third section, we cluster participants on the basis of their total number of vicarious touch endorsements and the number of reported exteroceptive and interoceptive sensations. We then show that the resulting clusters systematically vary on a number of other dimensions. Together, this analysis reveals several distinct vicarious touch profiles in our sample.

All statistical analyses in our study are Bayesian, including t-tests, ANOVAs, correlation assessments, and contingency tables. These were conducted using the R programming language ^53^. We employ the Bayes Factor R package to calculate Bayes Factors (BF) using the default prior and a Cauchy distribution centred at zero with scale parameter *r* = 0.707, which allows for a wide range of possible effect sizes ^54^. Bayes factors enable us to quantify the evidence for our alternative hypothesis against our null hypothesis. For instance, a Bayes factor of 3 indicates that the evidence provided by our data in favour of the alternative hypothesis is three times stronger than that for the null hypothesis. Generally, a Bayes factor > 1 provides evidence for the alternative hypothesis and a Bayes factor < 1 for the null, with Bayes factors between 1 and 3 often considered insufficient evidence in either direction ^55^. This Bayesian approach assesses the strength of evidence for one hypothesis over another without relying on dichotomous decision making, thereby permitting a more exploratory analysis without the pitfalls inherent to multiple comparisons in a frequentist framework.

## Results

### Broad characteristics

#### Prevalence & Intensity

We tallied the number of trials depicting touch on a hand out of 40 total where participants answered ‘yes’ to the question “*While watching the video, did you experience a tactile sensation on YOUR body?*”. Across all participants, 84% reported at least one such instance, leaving 16% never reporting it (Fig. 1). For those reporting vicarious touch, the average number of endorsed trials was 12.72 and the average intensity of these vicarious sensations was 3.44 (“*How intense was it?”* answer: 1 = not at all, 10 = extremely). Out of the whole sample, 70% of participants reported at least one *exteroceptive* sensation (a *tactile* sensation felt on the skin, including touch, pressure, scratching), which may be a stronger indication of sensory simulation than other vicarious sensations ^10^, and 50% of the whole sample reported vicarious *pain*. The relative frequency of all reported sensations was as follows: tingling (23%), pressure (18%), pain (15%), ticklish (13%), touch (11%), cold (6%), warm (6%), scratching (5%), and other (3%). Although ‘other’ responses were infrequent, an analysis of the free-text entries revealed that most participants provided specific tactile descriptors closely matching the observed touch (e.g., *soft, pinching, sharp, stabbing, tapping, stroking, pointy*; 79% of ‘other’ responses), while a smaller subset reported affective descriptors (e.g., *cringing, unease, tensing, shivering, discomfort*; 21% of ‘other’ responses).

**Figure 1.**
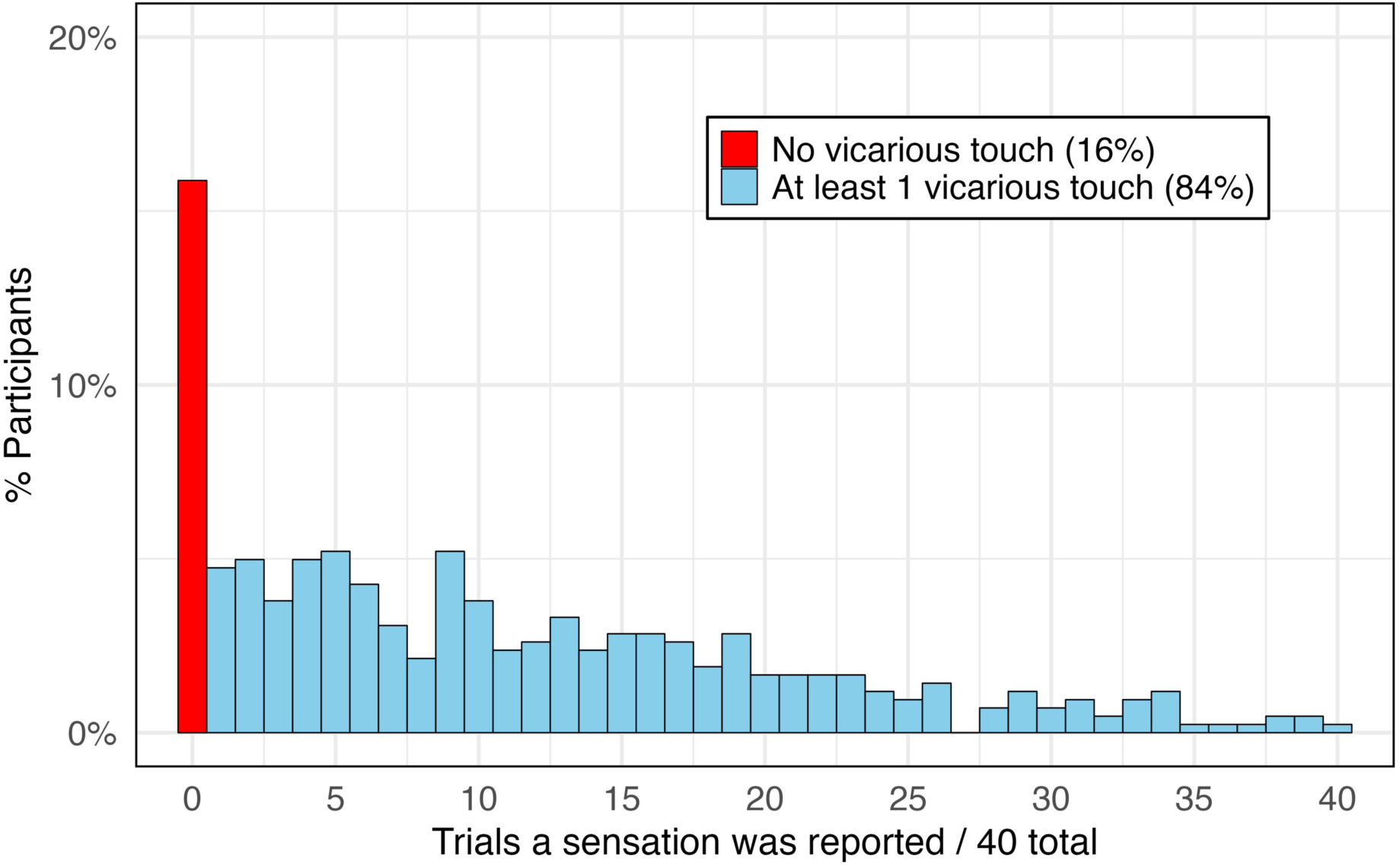
Distribution of the number of endorsed vicarious touch trials in our sample. The bar graphs represent the number of trials (out of 40 total) where participants (N = 422) reported a vicarious sensation after viewing 40 unique videos of a hand being touched. Each bar indicates the percentage of participants who reported the sensation across the specific number of trials (videos). Red indicates the percentage or participants who never reported vicarious touch, while blue represents those who reported it in at least one trial.

#### Location of vicarious touch

When participants reported experiencing a sensation while watching a video (all showing a right hand touching a left hand), we asked them where they felt it. We first computed the responses individually for each participant and then averaged these results across all participants for each body part. This approach ensures that our analysis reflects the typical participant response without being skewed by individuals with more frequent endorsements. Our findings indicate that, on average, 85% of trials where vicarious touch was endorsed involved the hands—53% in the left hand, 27% in the right hand, and 5% in both hands. Sensations in other body parts accounted for 11% of responses, 9% were described as a general feeling, and 0.7% were categorised as ‘other’ (these percentages exceed 100% because participants were allowed to select more than one option). Out of all participants who endorsed vicarious touch, 98% indicated a sensation in one or both hands in at least one trial. Thus, most vicarious touch responders experience sensations in a location congruent with the depicted tactile interaction—in this instance, the hand.

#### Gender differences

More women (88%) than men (76%) reported vicarious sensations in at least one trial, which was supported by a Bayesian contingency table (independent multinomial distribution with a prior scale parameter *a* = 1) indicating substantial evidence (*BF* = 7.67) against the null hypothesis of independence. An independent samples Bayesian t-test showed a higher average of 13.19 endorsed trials for women compared to 11.50 for men (*BF* = 4.40). The average intensity of these experiences did not clearly differ between the genders (3.52 for women vs. 3.25 for men: *BF* = 0.44). There was insufficient evidence regarding vicarious *pain* being more prevalent in women (54%) compared to men (41%) (*BF* = 2.32), although women did endorse feeling pain in a higher number of trials (4.82 vs. 3.64: *BF* = 9.5). There was again no clear difference in intensity (4.52 vs. 4.22: *BF* = 0.27). Finally, women had higher empathy scores than men in our sample (Women *M* = 11.22; Men *M* = 10.09, *BF* = 491.67)

#### Vicarious touch and emotional reactivity

To assess a link between vicarious touch and emotional reactivity, we calculated the Bayes factor for the Pearson correlation between the number of trials in which participants endorsed vicarious touch and their scores on the emotional reactivity subscale of the validated short EQ ^51^ (Fig. 2). There was inconclusive evidence regarding the small positive correlation between the number of trials in which participants reported vicarious touch and their empathy scores (*r* = 0.12, *BF* = 2.04). We also compared empathy scores between individuals reporting vicarious touch on at least one trial (EQ score *M* = 10.99) and those not reporting vicarious touch (EQ score *M* = 10.31). We found inconclusive evidence in the direction of the null (*BF* = 0.86) which specifies no difference in empathy scores between the two groups. These results were qualitatively the same for men and women (see Supplementary Fig. 2).

**Figure 2.**
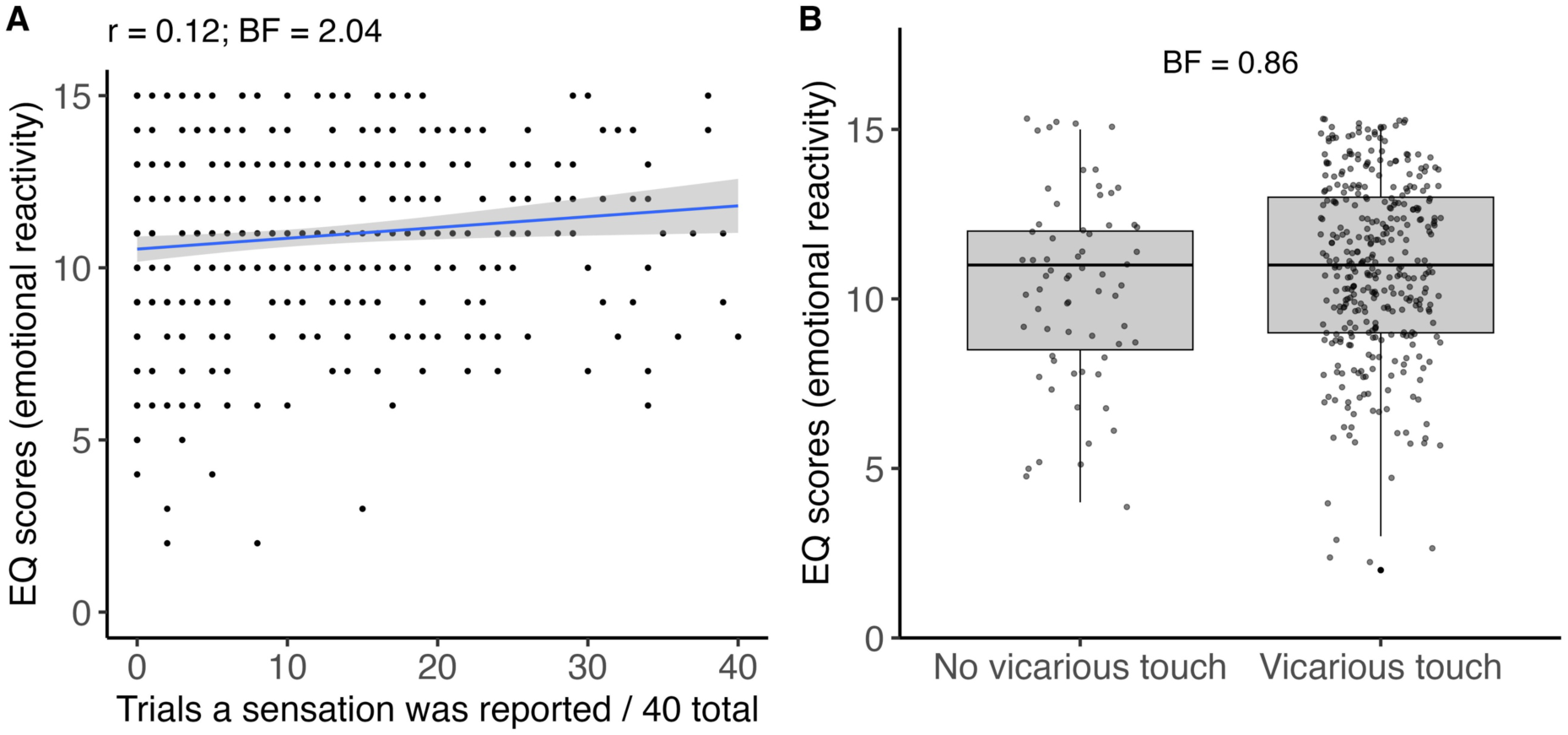
Vicarious touch endorsements and emotional reactivity scores. (Left) Scatterplot depicts the relationship between the number of vicarious touch endorsements out of 40 trials total and emotional reactivity scores based on five questions from the validated short EQ (higher scores suggest higher emotional reactivity). The line shows the linear regression with shaded 95% confidence intervals. (Right) Comparison between participants with no vicarious touch and at least one vicarious touch response. Scatterplots show the distribution of emotional reactivity scores for the two groups. The line within each box indicates the median, the box boundaries indicate the interquartile range, and the whiskers extend to 1.5 times the interquartile range.

### The influence of stimulus characteristics on vicarious touch experiences

#### Hedonic qualities of observed touch

For each individual video, we calculated the percentage of all participants who reported a vicarious touch sensation, and among those, we assessed the average intensity of the sensations. We then grouped the videos by their hedonic category (Fig. 3) as validated in the VTD ^15^. Based on prior research ^9,10,28^, we hypothesised that videos portraying pain or threat would be particularly effective in eliciting vicarious touch responses.

**Figure 3.**
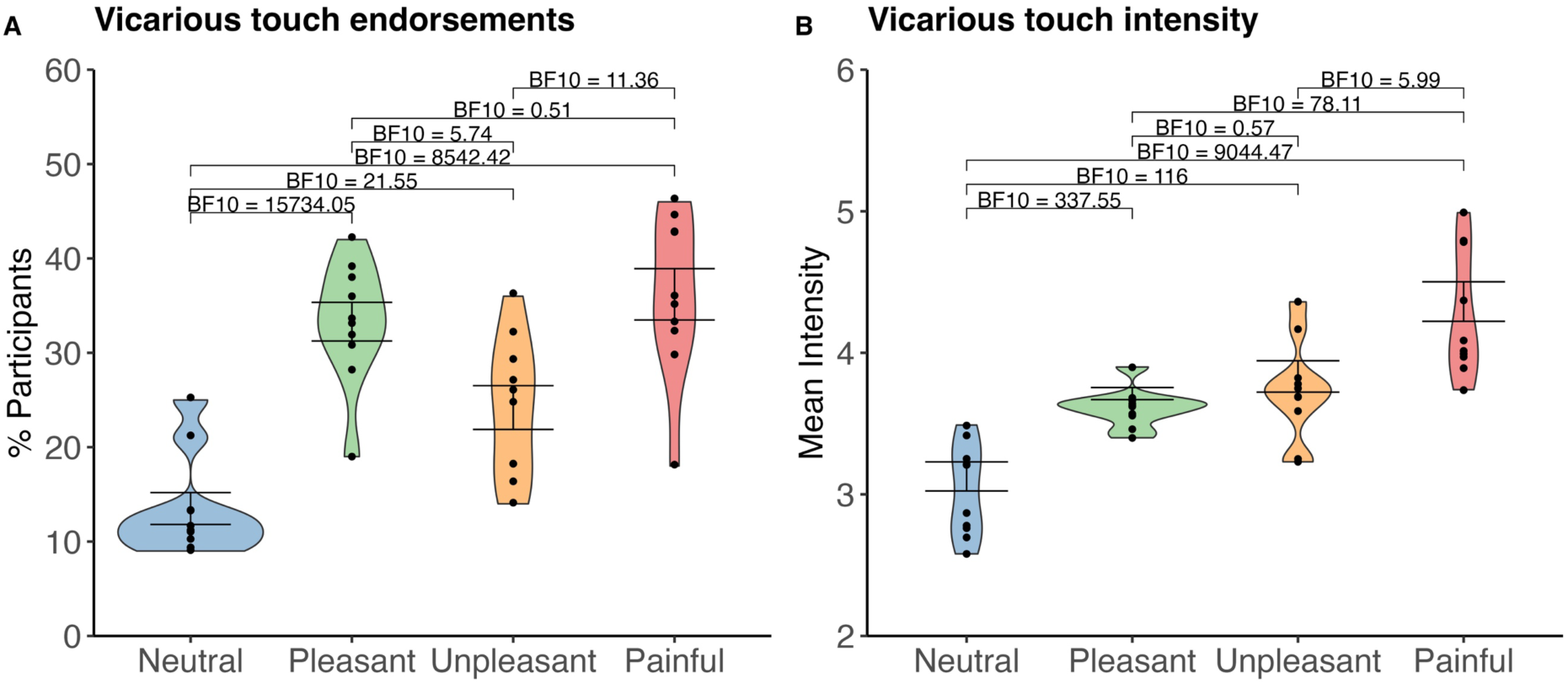
Vicarious touch endorsement and intensity across different hedonic categories. Dots (10 per hedonic category) show results for the individual videos. (A) Percentage of participants who endorsed vicarious touch for each video within the different categories. (B) Mean reported vicarious touch intensity for videos within the different categories (scale from 1-10; 1 = not at all, 10 = extremely). Violin plots convey the data distribution for each hedonic category: neutral, pleasant, unpleasant, and painful. The width of the violins indicates the density of the data, with broader sections representing higher densities. Error bars represent the 95% confidence intervals.

A one-way Bayesian ANOVA on vicarious touch endorsements provided strong support (*BF* = 313802.5) for a main effect of hedonic category. Post-hoc contrasts confirmed that painful videos were most frequently endorsed (36% of participants on average), with evidence for endorsement rates being higher than those for unpleasant (24%; *BF* = 11.36) and neutral videos (13%; *BF* = 8542.42) but not pleasant videos (33%; *BF* = 0.51). Similarly, pleasant videos were endorsed more frequently than neutral (*BF* = 15734.05) and unpleasant videos (*BF* = 5.74). The endorsement rate for unpleasant videos was higher than that for neutral videos (*BF* = 21.55). In sum, on average, painful and pleasant videos evoked vicarious touch most often, followed by unpleasant, and then neutral videos.

An analogous ANOVA on vicarious touch intensity also showed strong evidence for a main effect of hedonic category (*BF* = 2.34 x 10^39^). Post-hoc contrasts showed that painful videos evoked the highest average ratings (*M* = 4.26), relative to unpleasant (*M* = 3.73; *BF* = 5.99), pleasant (*M* = 3.61; *BF* = 78.11), and neutral videos (*M* = 3.03; *BF* = 9044.47). Unpleasant and pleasant videos were also rated higher than neutral videos (*BF* = 116.00 and *BF* = 337.55 respectively), but there was no clear evidence for difference between unpleasant and pleasant videos (*BF* = 0.57). Overall, on average, painful videos evoked the most intense sensations, followed by pleasant and unpleasant videos, and finally neutral videos.

In summary, participants indeed reported most frequent and intense vicarious sensations for painful videos, with neutral videos least likely to elicit vicarious touch and producing the lowest intensity when they did. These findings resonate with previous research ^9,10,28^ and underscore the impact of the hedonic quality of observed touch on the frequency and intensity of vicarious touch experiences.

#### The effects of arousal and threat on vicarious touch

Validation data from the VTD indicates that videos depicting painful touch are perceived as highly threatening and arousing ^15^, which might explain why they often elicit frequent and intense vicarious touch sensations. To examine this, we correlated the mean subjective arousal and threat ratings for each video (using the validation data) with the percentage of all participants that endorsed vicarious touch for each video and the mean intensity scores (Fig. 4).

**Figure 4.**
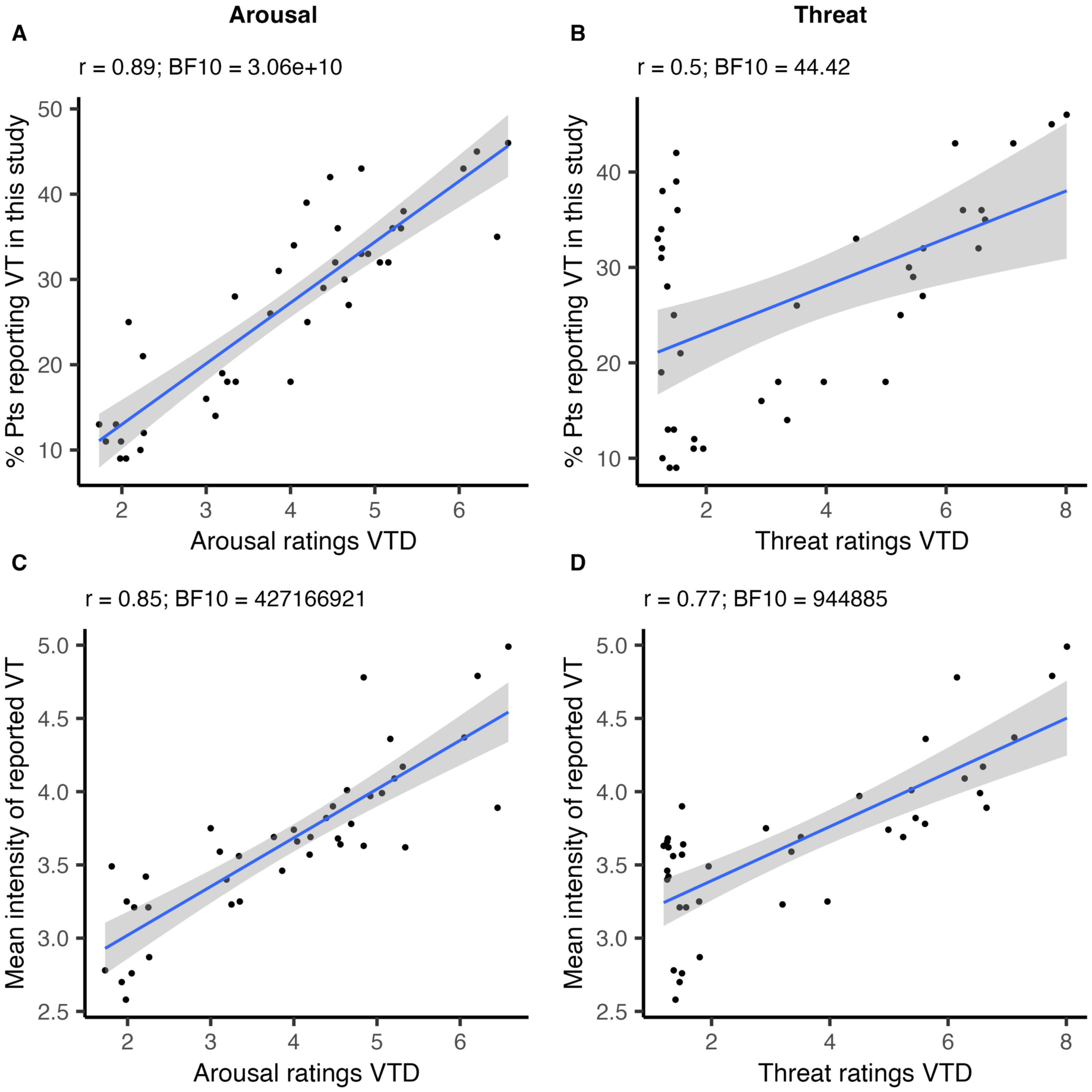
Correlation between the subjective arousal and threat ratings from the VTD and percentage of participants who endorsed vicarious touch (top row) and mean intensity (bottom row) for each video. Scatterplots depict the relationship between the independently rated (A, C) arousal and (B, D) threat on a scale from 1-10 (1 = not at all, 10 = extremely) from the VTD ^15^ and the endorsement (top) and intensity (bottom) of evoked vicarious touch for each video. Each dot signifies the percentage of participants that endorsed vicarious touch, or mean intensity ratings across participants, for the individual videos, with 40 videos in total. Lines show linear regression with shaded 95% confidence intervals.

The rated subjective arousal of the touch in videos was strongly correlated with both the percentage of participants who endorsed vicarious touch (*r* = 0.89, *BF* = 3.06 x 10^10^) and the average intensity of reported vicarious sensations (*r* = 0.85, *BF* = 4.27 x 10^8^). Threat ratings showed a moderate correlation with the percentage of participants who endorsed vicarious touch (*r* = 0.50, *BF* = 44.4) and a stronger correlation with the mean intensity of these sensations (*r* = 0.77, *BF* = 944885). However, closer inspection of Figure 4B suggests that these correlations are likely reduced by the cluster of observations with low VTD threat ratings but high vicarious touch endorsements. As shown in Supplementary Fig. 1 these observations derive entirely from neutral and pleasant videos. When correlations are computed separately on data grouped by hedonic video type, we observe strong correlations with unpleasant and painful videos for both vicarious touch endorsements (*r* = 0.89, *BF* = 20.05 and *r* = 0.78, *BF* = 5.11 respectively) and intensity (*r* = 0.69, *BF* = 2.89 and *r* = 0.72, *BF* = 3.57 respectively) but not for neutral and pleasant videos for vicarious touch endorsements (*r* = −0.07, *BF* = 0.63 and *r* = 0.54, *BF* = 1.47 respectively) and intensity (*r* = 0.37, *BF* = 0.89 and *r* = 0.42, *BF* = 1 respectively). A likely explanation for these findings is the limited range of threat ratings for neutral (1.27 – 1.95) and pleasant (1.19 – 1.52) videos, compared to the much broader range for unpleasant (2.92 – 6.59) and painful (4.5 – 8.01) videos. Additionally, there was a strong correlation between threat and subjective arousal ratings (*r* = 0.72, *BF* = 62473.23), indicating that videos perceived as more threatening also tend to be perceived as more arousing. These results collectively suggest that both rated arousal and threat are significant modulators that influence the likelihood and intensity of evoking vicarious touch sensations.

#### Sensations are felt across the body with a focus on the hands

We examined the specificity of sensation localisation in relation to the four hedonic categories used in the VTD (Fig. 5). Participants were asked the question, “Where on YOUR body did you feel the tactile sensation?” and could select multiple response options. We first calculated the average percentage per participant and then averaged these percentages across participants for each body part. This analysis revealed that the most common response was the left hand (video type: neutral 61%, pleasant 52%, unpleasant 52%, painful 51%), followed by the right hand (video type: neutral 30%, pleasant 25%, unpleasant 30%, painful 28%), and then both hands (video type: neutral 3%, pleasant 7%, unpleasant 7%, painful 5%). On some trials, participants indicated having a ‘general feeling’, which was more pronounced in response to observing painful videos (video type: neutral 1%, pleasant 7%, unpleasant 6%, painful 11%). In addition, we examined the responses described when participants indicated feeling touch on ‘another body part’ (video type: neutral 2%, pleasant 5%, unpleasant 3%, painful 4%). These reported sensations were dispersed across the body, especially in response to painful videos, with a notable concentration in the upper body regions, including the back, neck, chest, stomach, and arms. Participants rarely chose the ‘other’ option (neutral 0%, pleasant 1%, unpleasant 1%, painful 1%).

**Figure 5.**
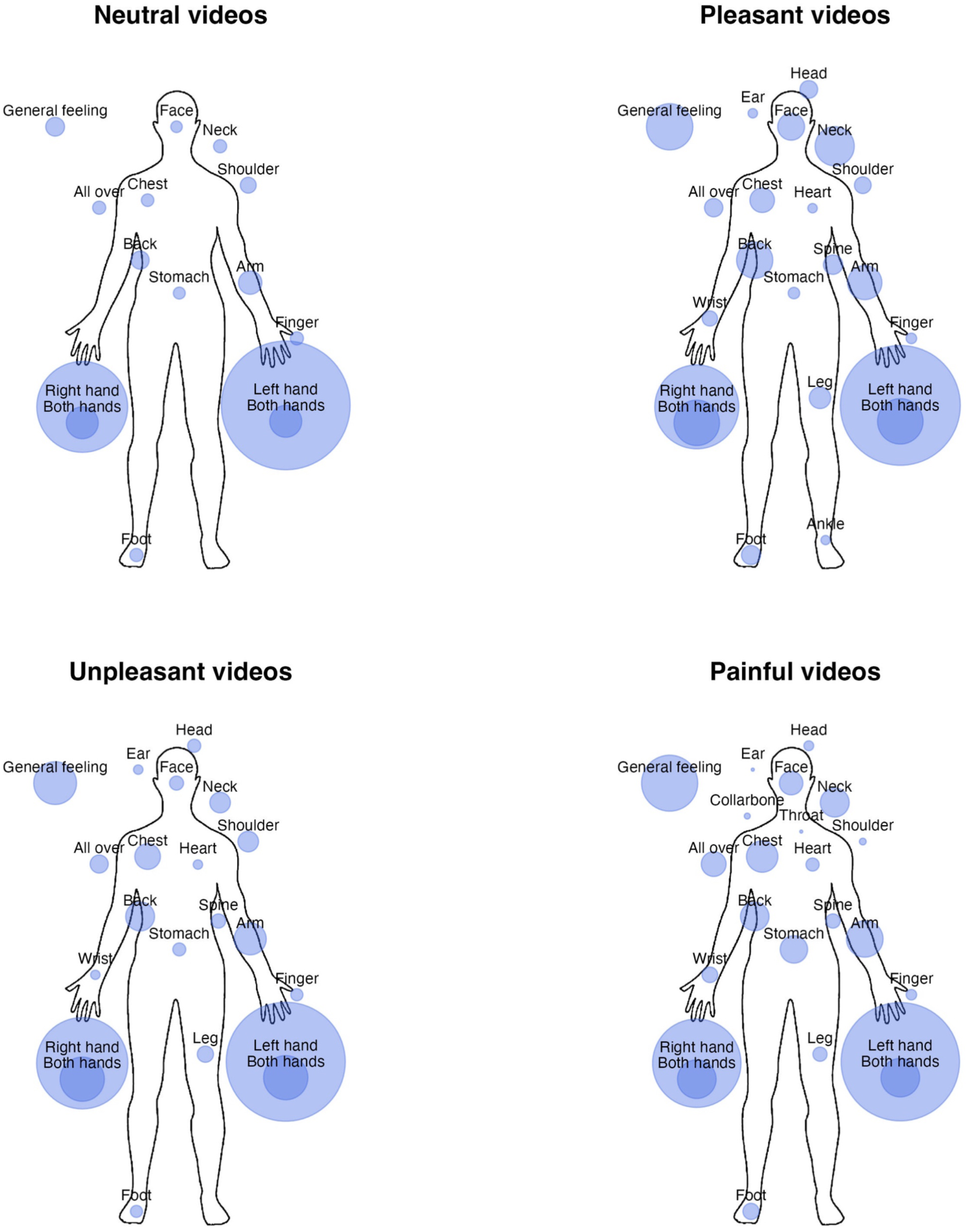
Location of reported sensations across different hedonic categories. Each body plot represents vicarious touch responses to 10 videos belonging to one of four hedonic categories: neutral, pleasant, unpleasant, and painful. Note that all videos depict a right hand touching a left hand. The response options were: Left hand, Right hand, Another body part (describe), A general feeling (describe), Other (describe). The body regions where sensations were felt are represented by purple circles. The size of these circles indicates the relative number of trials in which a sensation was reported in that region. For the hands, the larger lighter circle shows the responses for each hand separately with the darker blue showing responses of ‘both hands’.

#### Hedonic quality of vicarious touch response

In each trial where a vicarious sensation was reported, participants classified this vicarious sensation as being neutral, pleasant, unpleasant, or painful. We tested the general hedonic quality of the reported sensations (Fig. 6A), and the hedonic intensities (e.g., *how* pleasant was the touch?) (Fig. 6B). Across all participants, vicarious touch trials were most often categorised as ‘neutral’ (43%), followed by ‘unpleasant’ (34%), ‘pleasant’ (18%), and ‘painful’ (5%). A Bayesian ANOVA indicated that hedonic intensity scores were different across the four categories (*BF* = 7.72 x 10^23^). ‘Painful’ sensations were rated as most intense (*M* = 6.40), and this differed from both ‘pleasant’ (*M* = 5.11; *BF* = 8.21 x 10^9^) and ‘unpleasant’ (*M* = 4.63; *BF* = 2.13 x 10^18^) sensations, with additional evidence for a higher intensity for ‘pleasant’ compared to ‘unpleasant’ sensations (*BF* = 150.88).

**Figure 6.**
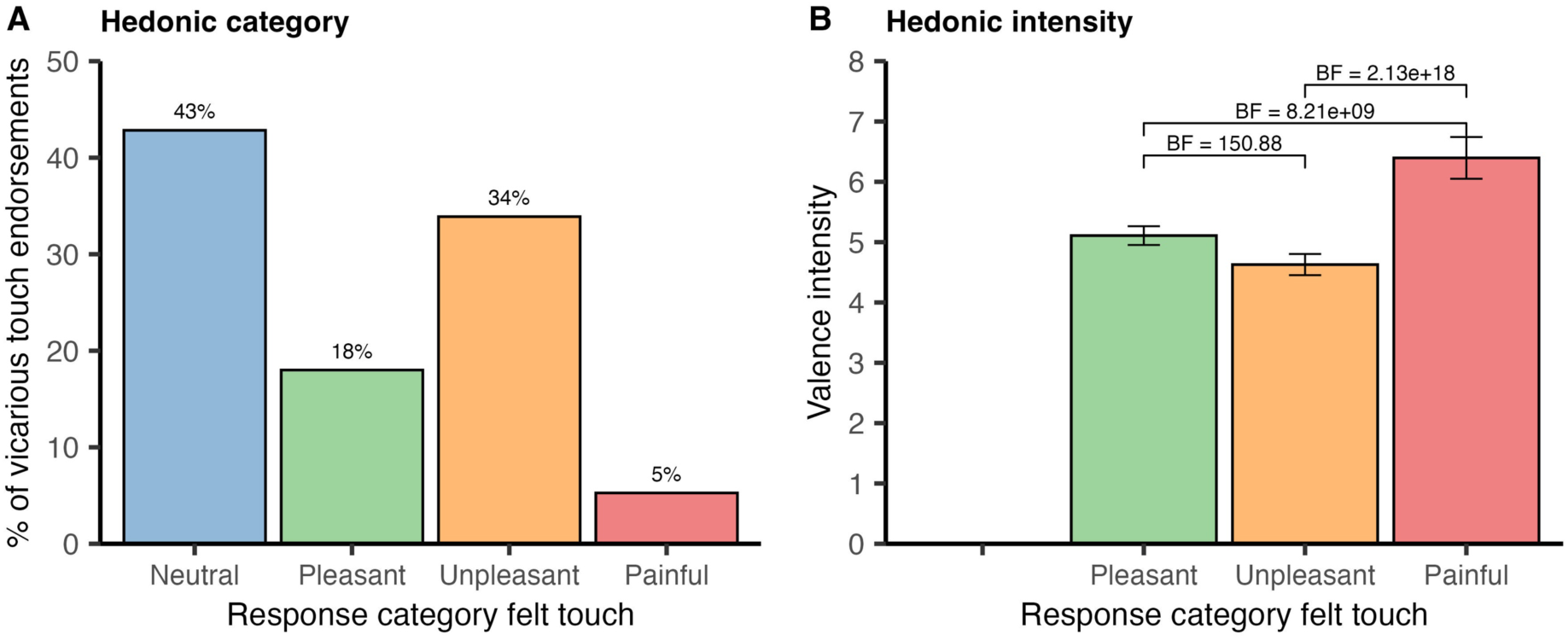
Vicarious touch responses with different hedonic qualities. (A) The percentage of all endorsed vicarious touch trials in which participants categorised the felt touch as neutral, pleasant, unpleasant, or painful. (B) The corresponding average intensities for each hedonic quality (participants were only asked about hedonic intensity – how pleasant/unpleasant/painful was the touch? - for non-neutral responses). Intensity was scored on a scale from 1-10 (1 = not at all, 10 = extremely), with higher values indicating stronger sensations. Error bars represent 95% confidence intervals for each category.

#### The hedonic quality of felt touch sensations

Next, we tested whether the hedonic quality of the observed touch portrayed in the videos influences the hedonic quality of reported sensations (Fig. 7). Neutral videos predominantly evoked neutral sensations (78%), while both pleasant and unpleasant videos frequently elicited either hedonically-congruent sensations (48% and 45% respectively) or neutral ones (46% and 47% respectively). Notably, when participants viewed videos showing painful touch, they predominantly reported the sensation as being unpleasant (64%), with few instances of painful (12%). This implies that the hedonic quality is regularly simulated but in a weaker way for videos depicting painful touch.

**Figure 7.**
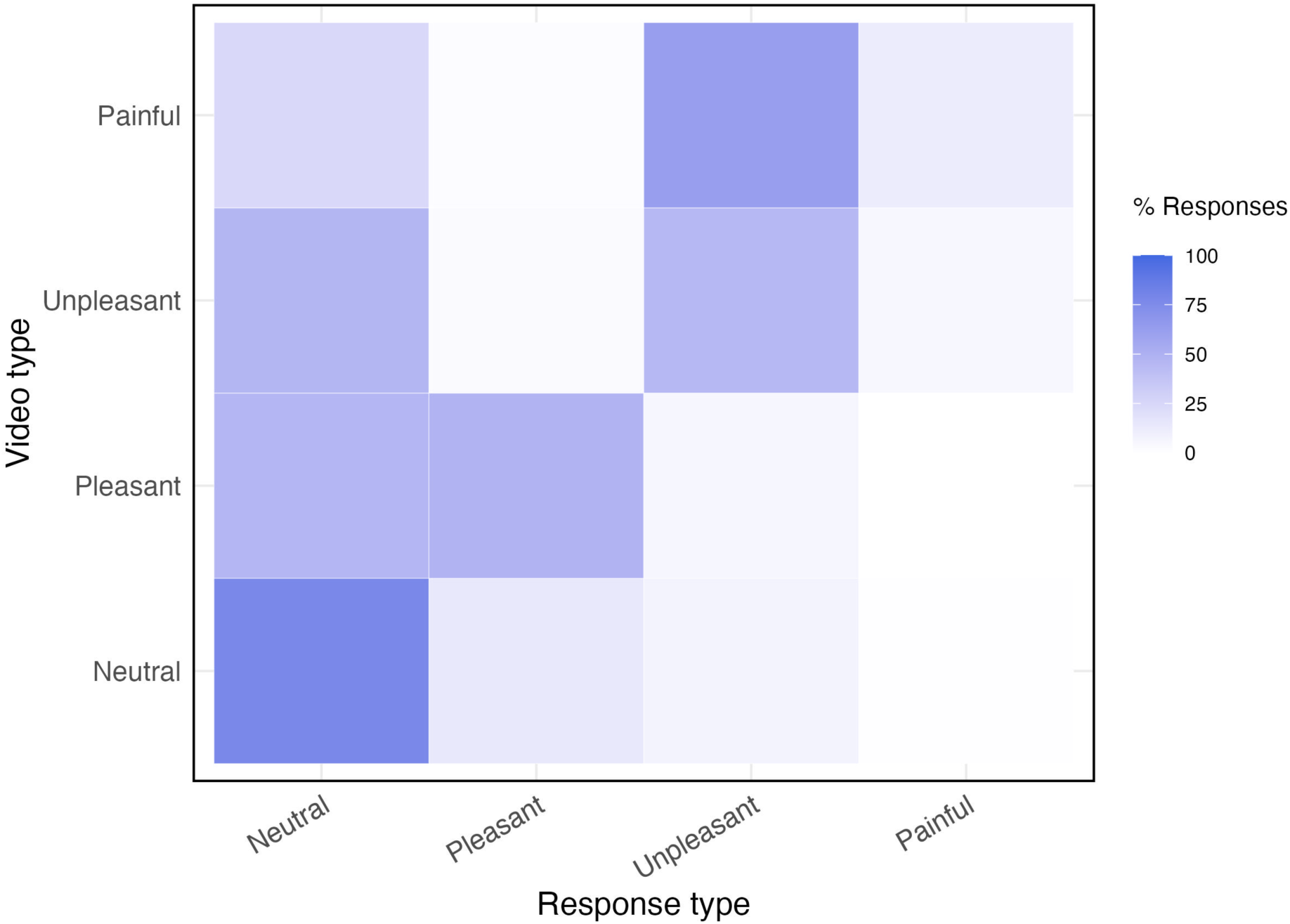
Impact of hedonic quality of video touch on vicarious touch sensations. The colour intensity represents the percentage of trials where sensations were reported with a hedonic quality for each of the ten videos within each category. Hedonic categories of the videos are predefined based on validation data from the VTD ^15^.

To further explore the relationship between the quality of the observed and felt touch, we examined whether the hedonic intensity also matched. Participants first categorised their sensations as neutral, pleasant, unpleasant, or painful (i.e., see Q4 in the methods section), and then rated the hedonic intensity of *the felt touch* (e.g., if they indicated that the sensation was pleasant, we would ask *how* pleasant on a scale from 1-10; see Q5 in the methods section). These ratings were compared to scores from the VTD ^15^ where an independent cohort had rated the hedonic intensity of *touch in the videos* using the same scale. Our analysis revealed that on average, within each hedonic category, the mean ratings obtained across videos in from the VTD was very similar to the mean ratings of vicarious touch reported here. In particular, the average VTD rating for pleasant touch was 6.26, with vicarious sensations averaging 5.11; the average VTD rating for unpleasant touch was rated at 5.15 with vicarious sensations averaging 4.63; the average VTD rating for painful touch was 7.11, with vicarious sensations at 6.40. Overall, the hedonic intensity of felt touch was consistently rated slightly lower than that of observed touch across all categories (1.15, 0.52, and 0.71 lower for pleasant, unpleasant, and painful, respectively). Upon further inspection, however, we found that, while there was a clear relationship in *average* ratings across hedonic categories as just described, there was noticeable variability in hedonic matching for the individual videos in each category (see Supplementary Fig. 5). Overall, we found that 1) reported vicarious touch experiences often have an hedonic quality, 2) the *average* rated hedonic intensity is very similar for both observed touch in the videos and vicarious touch felt on the body, but slightly lower than that of observed touch across all categories, and 3) there was variability in how well the seen and felt touch matched in hedonic intensity when considering the individual videos.

#### Different types of vicarious sensations

We investigated the type of sensation elicited by viewing touch videos of different hedonic qualities, such as whether the participants described tingling, touch, temperature, pressure and so on. To achieve this, we calculated the percentage of each type of reported sensation relative to all reported sensations across the ten videos within each of the four categories (Fig. 8).

**Figure 8.**
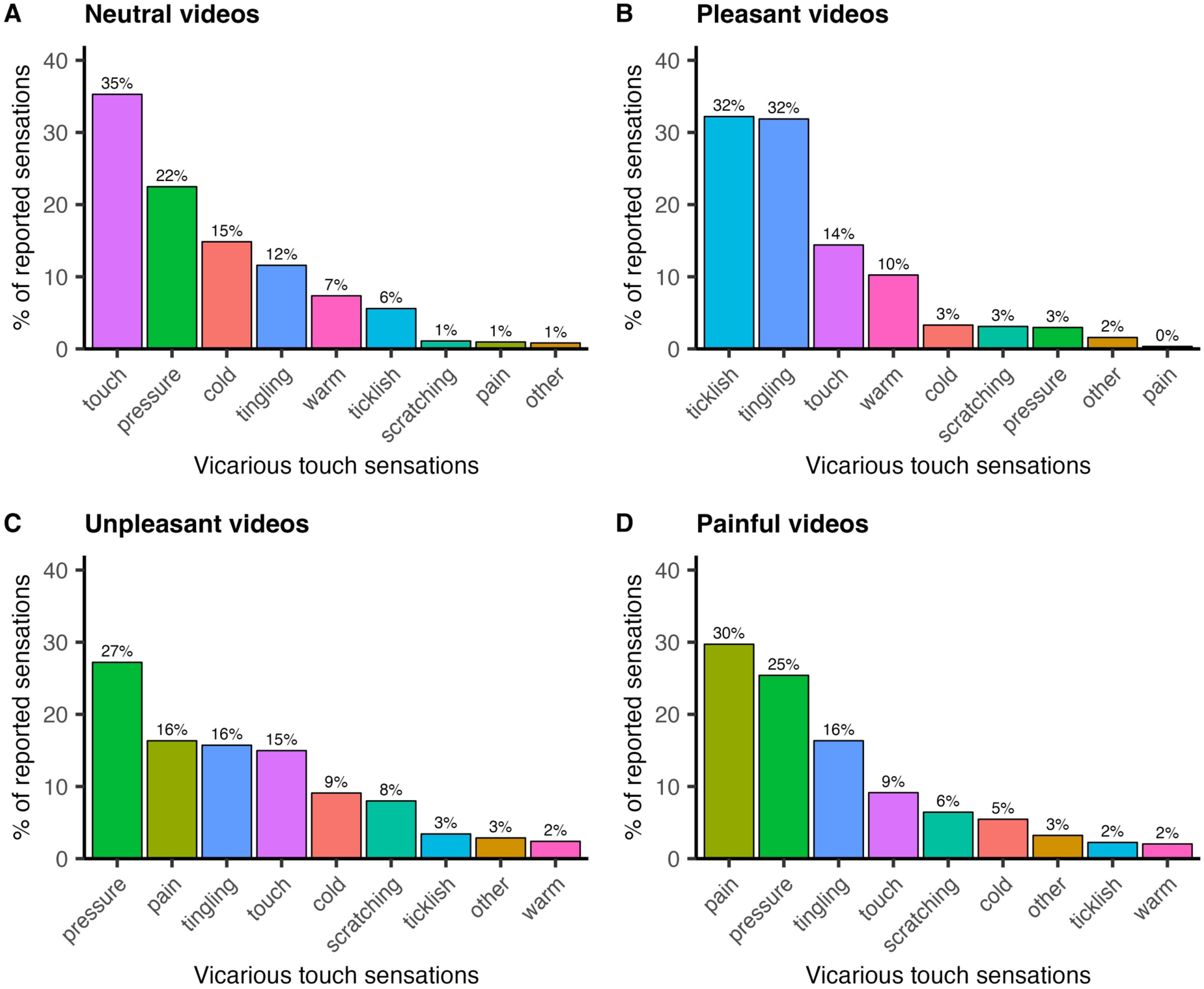
Differential sensation reporting across hedonic video categories. The distribution of vicarious touch sensations reported by participants following the viewing of ten videos categorised as (A) Neutral, (B) Pleasant, (C) Unpleasant, and (D) Painful. The y-axis displays the percentage of each sensation type relative to all reported sensations across the ten videos within each of the four categories. The colour of the bars code for sensation type and this is consistent across all four panels.

Our findings revealed that the quality of the touch depicted in the videos tended to produce vicarious sensations with similar qualities (see Supplementary Fig. 3 for sensation reporting for each of the 40 videos). Specific mirroring occurred on a fine-grained level: for example, participants reported feeling mostly pressure when touch with scissors was applied forcefully from the top to the bottom of the hand, but they reported mostly tingling when a similar touch with scissors was applied softly. Overall, neutral videos, often depicting a singular touch application with moderate pressure ^15^, primarily elicited sensations of ‘touch’ and ‘pressure’. Pleasant videos, showing a gentle stroke using soft objects like brushes, fabrics, or another hand, predominantly evoked ‘ticklish’ and ‘tingling’ sensations. Unpleasant videos, where the touch was generally applied with significant pressure, triggered responses of ‘pressure’ and ‘pain’. In videos categorised as painful, which often portrayed touch using strong pressure or a sharp object, the predominant vicarious sensations were ‘pain’, and ‘pressure’. Videos categorised as pleasant sometimes evoked warm sensations, while those deemed unpleasant more often led to cold sensations, likely reflecting the temperature-related properties of the objects used, such as soft materials or a flat hand (warm) versus a metal spoon or knife (cold). These findings suggest that the type of touch, the intensity, pressure, and the characteristics of the object being used to perform the touch, shape the specific sensations experienced.

#### Observed touch on an object

Previous research suggests that although observing touch on either humans or objects can elicit vicarious touch sensations, these responses are more common and intense for touch on humans ^10,28^. Similarly, in our study, we found that only 32% of all participants reported at least one vicarious touch when observing touch on an object (5 trials total), compared to the much higher 84% who had at least one vicarious touch instance when observing touch on a *hand* (40 trials total). This is not just because we had less object videos: on average, a single video showing touch on the hand evoked sensations in 27% of participants, whereas touch on the object or adjacent to the object elicited responses in only 12% and 7% of participants, respectively. The type of touch on the object influenced vicarious responses: a soft brush stroke on an object triggered sensations in 22% of participants, nearly three times as much when the object was simply touched with a finger which evoked sensations in 8% of participants. Participants described similar sensations in line with the type of touch (e.g., touch with a finger, Fig 9). These findings suggest that while touch observations to another human have a higher propensity to elicit vicarious sensations, the type of sensations reported are the same, irrespective of whether the target is human or an object.

**Figure 9.**
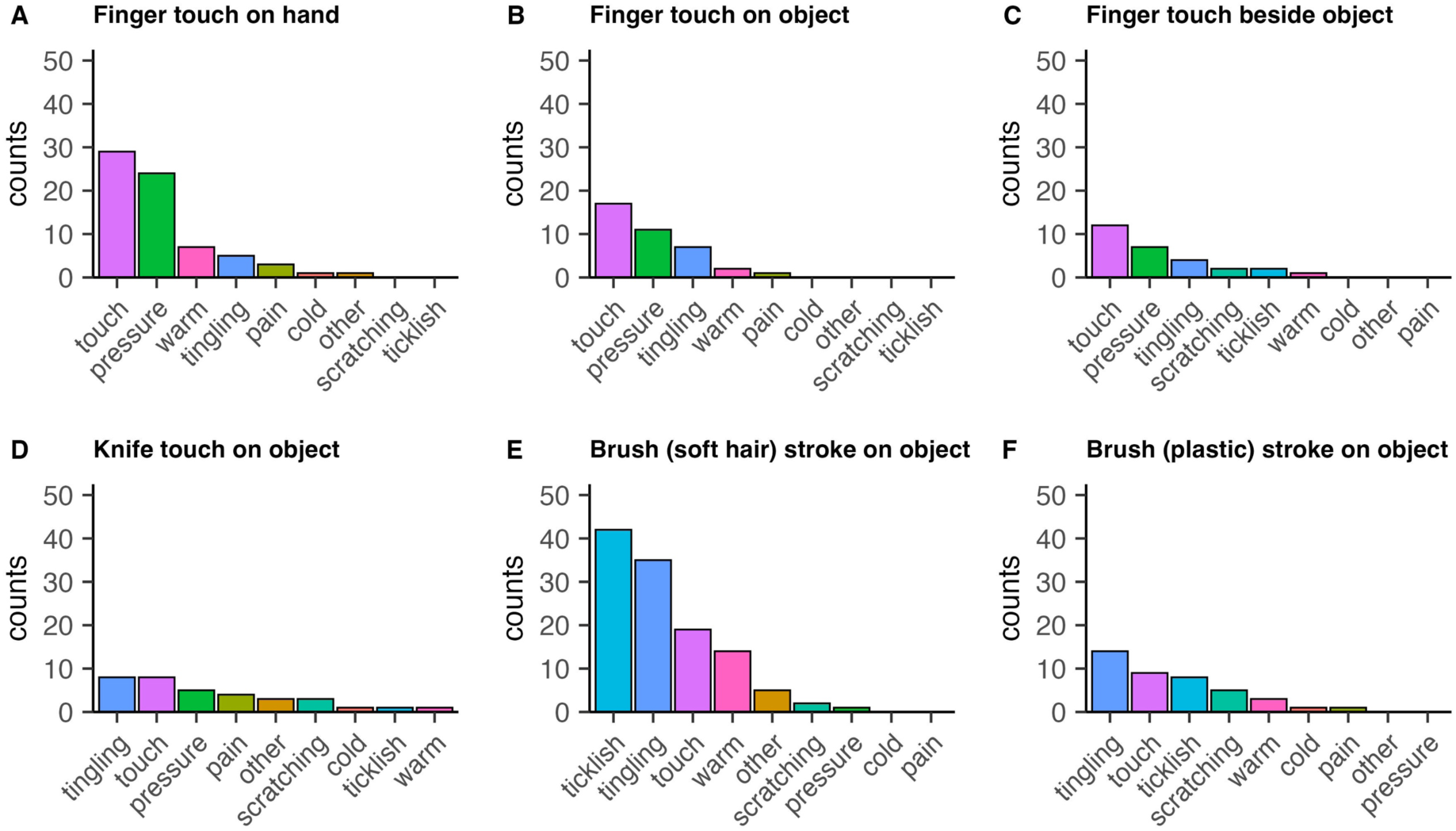
Sensations reported when observing touch on an object or beside an object. (A, B, C) Comparison of reported sensations for videos depicting identical touch interactions with a hand, an object, and beside an object. (D, E, F) Responses to the remaining object-touch videos (a total of 5 object videos). Note that the y-axis represents the *total counts* of reported sensations rather than relative frequencies to illustrate the frequency of sensations for hand-touch versus object-touch. Sensations are ordered by the highest count.

We next investigated the locations where participants reported feeling touch while observing the object videos. We again established the responses for each participant and then averaged across participants. Participants could report sensations in multiple areas for each trial. The reported locations were: **right hand** (29%), **left hand** (28%), **general feeling** (12%), **both hands** (6%), **face** (2%), **back** (2%), **neck** (2%), **arm** (2%), **chest** (1%), **stomach** (1%), **finger** (1%), **leg** (1%), **spine** (<1%), **all over** (<1%), **ear** (<1%), **head** (<1%). The predominance of sensations in the hands suggests a possible priming effect from the other videos, which featured touch on a left hand, or a general tendency among participants to mirror observed touch on objects onto their own hands. One obvious explanation is that when a right index finger is the touching agent, this may result in evoked touch (see also ^10^). We therefore analysed responses specifically from the two videos where a right-hand finger was the touching agent. Here, reported sensations were indeed most likely to be localised to the **right hand** (48%), with other reports of **left hand** (27%), **both hands** (6%), **general feeling** (3%), **finger** (1%), **head** (<1%), **chest** (<1%), **stomach** (<1%), and **leg** (<1%). Overall, we found that while the sensations reported are similar regardless of the target, observed touch to a human evoked a much higher prevalence of vicarious sensations than to an object.

### Different vicarious touch profiles in a general population

In this third section of our findings, our objective was to determine if our data revealed subgroups of individuals who experience vicarious touch in qualitatively distinct ways. To test this, we applied a clustering approach based on the total number of trials in which a participant: 1) endorsed vicarious touch (i.e., answered ‘yes’ to the first question), 2) reported an exteroceptive sensation, or 3) reported an interoceptive sensation. For example, if in a trial where vicarious touch was endorsed, a participant reported two exteroceptive sensations which are felt on the skin (e.g., touch and pressure) and one interoceptive sensation which is felt from within (e.g., tingling), this would count as one instance for each of the three categories, regardless of the number of sensations reported. We only included vicarious touch responders, i.e., participants who experienced at least one instance of vicarious touch. Drawing on prior research ^10^ we hypothesised that individuals with a higher number of total vicarious touch endorsements would report relatively more exteroceptive sensations compared to those with fewer vicarious touch endorsements, who may experience relatively more interoceptive sensations. This hypothesis is based on the idea that stronger vicarious responses may more closely mirror the sensory qualities of the observed touch, whereas weaker vicarious responses may be experienced in a more diffuse or internalised manner ^3,8,10,19^. It is important to note that some sensations, such as tingling and tickling, may have ambiguous origins and could also arise from externally perceived stimuli ^19^ (see the Discussion section). Our categorisation aimed to reflect broad differences in sensory qualities rather than strict dissociations.

We used k-means clustering and the Elbow Method ^56^ to determine the optimal number of clusters for our dataset (see Supplementary Fig. 4). We analysed the difference between successive values of the within-cluster sum of squares (WCSS) ^57^ and used the point of inflection to select our number of clusters. This showed that two or three clusters would be the most appropriate number of clusters based on the variance explained. We decided to include three (rather than two) clusters to increase the likelihood we would capture those on the more extreme ends of the vicarious touch distribution.

We assessed compactness within each cluster by calculating the average of all pairwise Euclidean distances among observations, reflecting how closely data points are grouped. In addition, inter-cluster distances were calculated as the average of all distances between points across different clusters, providing insight into the distinctness and separation of each cluster. These measurements demonstrated differentiation between clusters, with intra-cluster distances averaging 8.67 (range = 5.53 to 12.04) and inter-cluster distances averaging 23.74 (range = 15.31 to 34.18). The clustering validity was further supported by a Dunn Index ^58^ of 1.271, indicating reasonable separation and compactness. Additionally, the Silhouette Score ^59^, averaging 0.504, confirmed that the clusters are well-separated, aligning with the Dunn Index findings. This combination of metrics validates our clustering approach, demonstrating both effective separation and internal cohesion within clusters.

Figure 10 presents a scatter plot of the clusters, with the number of interoceptive sensations on the x-axis and the number of exteroceptive sensations on the y-axis. The size of the markers represents the total number of reported vicarious touches. Note that the total number of vicarious touches is not simply the sum of interoceptive and exteroceptive sensations, as participants could report both types of sensations in a single trial. Cluster 1 (purple), comprising 124 participants (36% of vicarious touch responders; 95 women, 28 men, 1 non-binary), displayed the lowest levels of vicarious touch endorsements (*M* = 5.56, range = 1-14) and reported more interoceptive than exteroceptive sensations (*M* = 3.18 vs. 2.09). This can be seen in Figure 10, where more Cluster 1 points are scattered below the diagonal line. Cluster 2 (green), the largest group with 187 participants (53% of vicarious touch responders; 129 women, 58 men), showed moderate vicarious touch endorsements (*M* = 17.11, range = 10-29) and relatively more interoceptive compared to exteroceptive sensations overall (*M* = 9.66 vs. 7.15). However, the distinction between the two sensation types was less pronounced compared to Cluster 1 as can be seen in Fig. 10. Cluster 3 (yellow), comprising 44 participants (12% of vicarious touch responders; 32 women, 12 men), registered the highest vicarious touch endorsements (*M* = 30.80, range = 22-40) and had a notably higher proportion of exteroceptive compared to interoceptive sensations (*M* = 21.07 vs. 13.93). In Figure 10, Cluster 3 points are predominantly located above the diagonal line.

**Figure 10.**
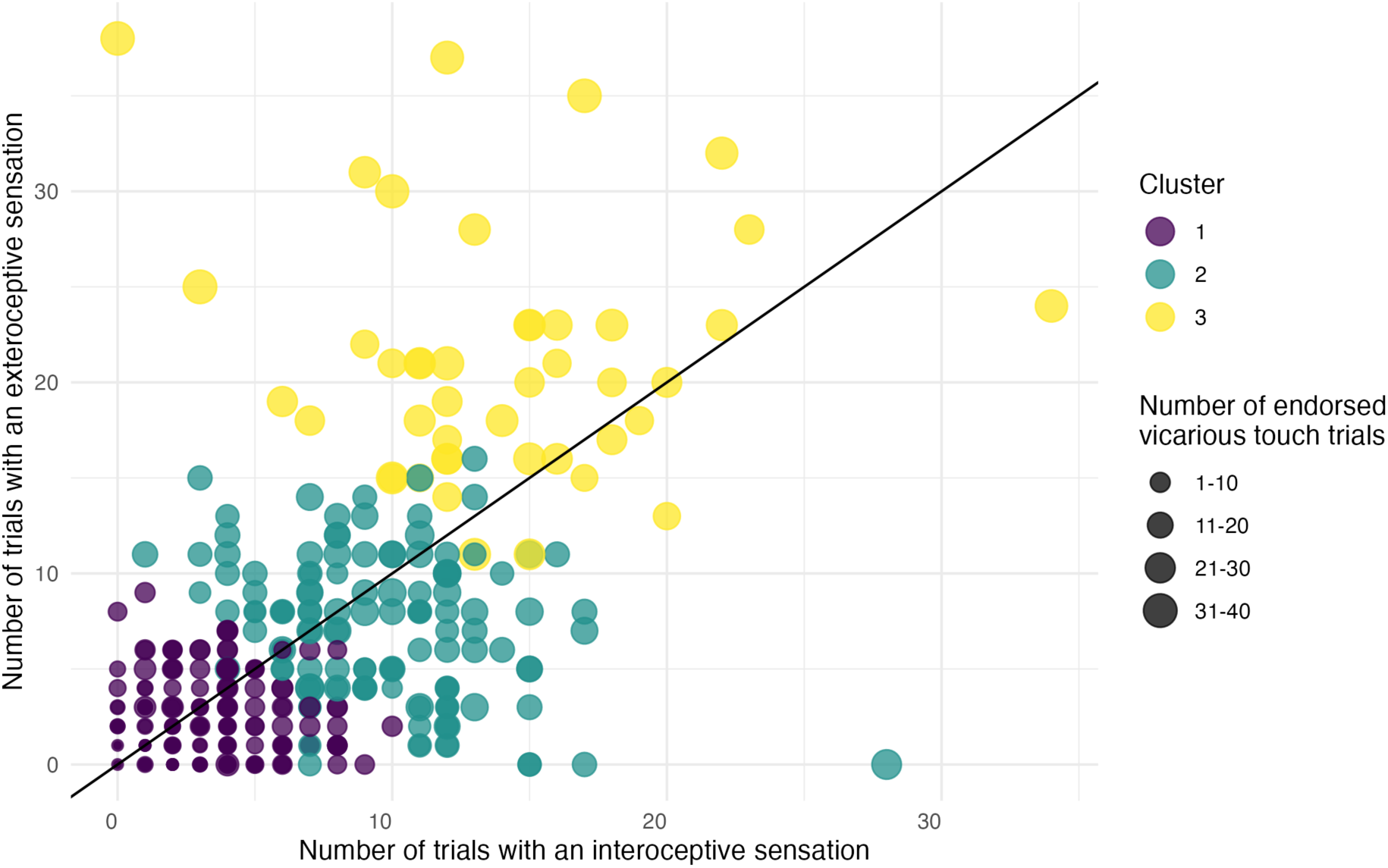
Scatter plot representing the results of a data-driven cluster analysis based on the number of trials with endorsed vicarious touch, exteroceptive sensations, and interoceptive sensations. Each point represents a single participant. The size of the points indicates the total number of endorsed vicarious touch trials, while the colour denotes cluster identity. The diagonal line indicates an equal number of interoceptive and exteroceptive sensations. This line helps distinguish the relative proportions of sensations for each cluster, with points above the line indicating more exteroceptive sensations and points below the line indicating more interoceptive sensations.

In addition to variations in the total number of vicarious touch endorsements and the interoceptive and exteroceptive sensations used for the clustering algorithm, these clusters exhibit additional distinguishing characteristics not used as inputs (see Fig. 11). Panel A displays the proportion of trials where reported sensations were localised to the hands for videos depicting touch on a hand, relative to the total number of trials where vicarious touch was endorsement. This panel reveals that individuals in Cluster 3 reported hand-localised vicarious touch more frequently than those in the other two clusters. Panel B, related to videos depicting touch on a hand, shows the proportion of trials with endorsed pain sensations relative to the total number of trials with endorsed vicarious touch. It indicates that Cluster 2 individuals reported feeling pain in a higher proportion of trials than those in Cluster 1, and similarly, Cluster 3 reported pain sensations in relatively more trials than Cluster 2. While the evidence supporting these two comparisons was limited, there was substantial evidence confirming the differences in pain sensations between Clusters 3 and 1. Panel C illustrates the total number of vicarious touch endorsements when observing touch on an object. In this case, Cluster 2 endorsed more trials than Cluster 1, and Cluster 3 endorsed more trials than Cluster 2, suggesting that broader underlying mechanisms such as arousal or attention may influence vicarious touch experiences. This variation could also reflect different thresholds or criteria for reporting vicarious touch across clusters.

**Figure 11.**
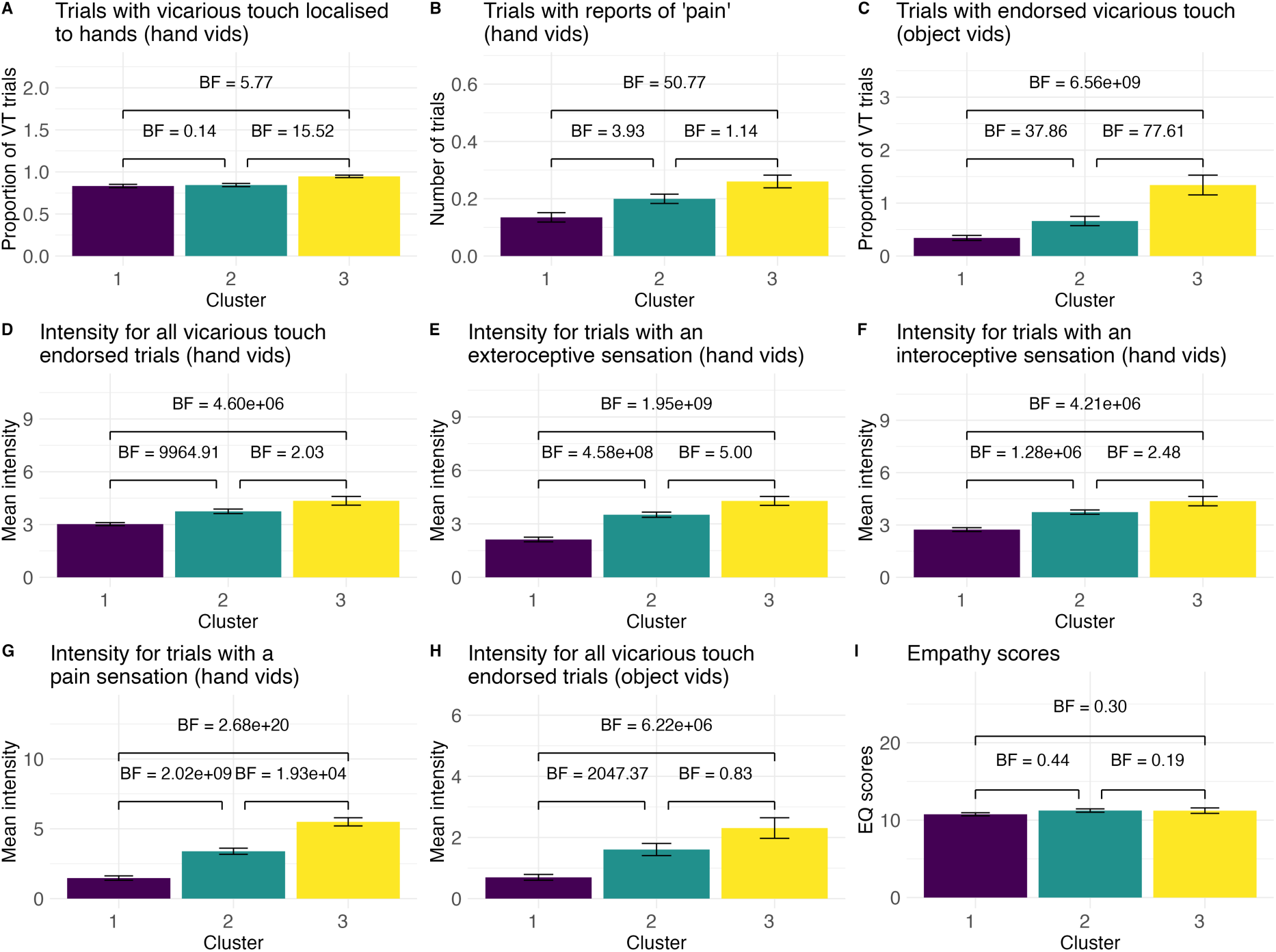
Bar plots showing variations in different response variables across each cluster. Panels A-C illustrate the relative proportion of specific sensations in response to certain videos, normalised by the total vicarious touch endorsements per cluster. Panels D-G display the mean intensity for different response types and H for observed touch on an object. Error bars indicate the standard error of the mean. Bayes Factors comparing the clusters are shown above each pair of bars, indicating the strength of evidence for differences between the clusters

In terms of intensity, the most notable contrast was between Clusters 1 and 2, with Cluster 2 showing higher intensity across all sensation types (panels D-G) and when observing touch on an object (panel H). This indicates that for those with a low to moderate tendency to report vicarious touch, an increasing tendency to report vicarious touch was associated with higher intensity overall. However, we found moderate to strong evidence of no differences among the clusters in distinguishing between observed touch on a person versus an object, as measured by the intensity for human versus object touch (Cluster 1 vs. Cluster 2 *BF* = 0.28, Cluster 2 vs. Cluster 3 *BF* = 0.19, and Cluster 1 vs. Cluster 3 *BF* = 0.18). Interestingly, individuals in Cluster 3 did not generally experience vicarious touch with greater intensity than those in Cluster 2, except for ‘pain’ (panel G), which was most intense for Cluster 3. Given the relative muting of pain sensations reported earlier, this result may suggest that mirroring pain is a special case compared to mirroring non-painful touch. The clusters did not differ in their emotional reactivity scores (panel I), suggesting that empathy for sensations may be dissociable from other types of empathy.

## Discussion

We investigated the characteristics of reported vicarious touch among a general population, using a controlled set of video stimuli from the Validated Touch-Video Database (VTD) ^15^ that depict various types of touch between hands. The findings revealed that a large majority of participants (84%) reported vicarious sensations such as tingling, pressure, and pain when watching videos depicting various types of touch. This high prevalence underscores a broad capacity for sensory mirroring in the general population that extends beyond the specialised phenomenon of mirror-touch synaesthesia.

### Prevalence

Our study revealed a prevalence of vicarious touch that exceeds prior studies, with 84% of participants reporting at least one sensation during trials that depicted touch on another person’s hand. For comparison, earlier papers reported that approximately 25-30% of a Western undergraduate sample ^10,16^ and 60% of a Chinese sample ^9^ similarly reported at least one vicarious sensation. In our sample, 70% of participants also specifically reported at least one *exteroceptive* sensation such as touch, pressure, or scratching, which is thought to be a stronger indicator of sensory mirroring than more general activations like tingling ^10^. Additionally, our study found that 50% of participants reported experiencing pain specifically when viewing videos depicting painful touch. This rate is higher than that of prior research that focused more on general depictions of pain ^9–14^. This implies that the specific portrayal of painful *touch* might activate more direct sensory pathways, leading to a higher incidence of localised pain sensations.

Several methodological factors may explain the high prevalence of vicarious touch in our study. Perhaps most importantly, our stimuli are close-up, dynamic videos of self-touch interactions, many of which were emotionally charged, likely enhancing engagement and drawing participants’ attention ^20,60,61^. Perspective may have also played a role ^38^, as the first-person viewpoint could have encouraged an embodied simulation of the observed hand ^37^ rather than, or in addition to, an empathic response to another person’s experience. These factors may have contributed to the increased reports of vicarious touch compared to prior research.

Additionally, self-report measures introduce the possibility of response bias ^62–65^, where participants may report expected experiences rather than genuine ones, potentially inflating vicarious touch rates. However, the specificity in ratings—such as stronger endorsements for painful versus neutral touch—suggests this bias did not overly skew results. Another factor is phenomenological control, where visual stimuli act as imaginative suggestions shaping subjective experiences ^66,66^. This may have promoted stronger vicarious touch responses in our experimental setting compared to real life ^66^.

Future research could complement subjective reports with objective measures, such as neuroimaging, to explore the neural mechanisms underlying vicarious touch. For example, our recent EEG study found that participants who reported vicarious touch from VTD videos exhibited neural overlap between observing touch and experiencing it directly, a pattern not seen in those without vicarious touch responses ^4^. This suggests a neural basis for individual differences in vicarious touch processing.

### Gender & Empathy

Although the overall intensity of reported sensations did not differ between genders, women more frequently reported vicarious sensations than men, both in the proportion experiencing at least one vicarious touch and the average number of endorsed trials. They also reported pain sensations more often. These findings contrast with previous research showing no gender differences in the perception of non-painful vicarious touch ^9,10^, but align with studies indicating that women are more likely than men to report pain when observing others in pain ^12,16^. This suggests that conscious vicarious touch experiences may be more prevalent in women, including a tendency to report pain-related sensations. Some neuroimaging studies have also shown that women show stronger activation in sensory and affective brain regions when observing others in pain ^43,44^, but see ^67^. Cultural norms may encourage women to be more attuned to social and emotional cues ^68,69^, potentially influencing their neural responses and self-reported vicarious sensations. However, as our study found no gender differences for intensity, it remains unclear whether these differences reflect a heightened physiological response, increased self-awareness, or different expectations, in reporting bodily experiences.

Women in our sample also scored higher on the emotional reactivity subscale of the short Empathy Quotient (EQ) ^51^, consistent with previous research showing higher self-reported empathy in women ^62,70–72^. However, we found did not find conclusive evidence for a correlation between vicarious touch and emotional reactivity scores, suggesting that heightened emotional reactivity likely did not drive the observed gender differences regarding vicarious sensation. This observed lack of correlation aligns with some findings ^4,45^ but contrasts with others ^10,29,46^. Our findings suggest that experiencing vicarious touch is not merely a generalised amplification of resonance with others across both sensory and emotional domains ^73^. Emotional reactivity involves largely automatic, affective responses to others’ emotions ^51,74^, whereas vicarious touch may depend more specifically on sensory processing and embodied simulation of observed touch. Understanding these differences could provide deeper insights into the multifaceted nature of empathy and inform interventions aimed at enhancing empathic abilities.

Our study used validated touch videos that were standardised in terms of hand appearance, placement, and perspective, controlling for variables that may influence vicarious touch responses ^28,33^. This methodological approach enhances the reliability of our findings and isolates the sensory aspects of touch. However, identification with the observed individual has been shown to modulate vicarious sensations ^75,76^. Despite lacking strong identity cues and being neutrally presented, our stimuli did all show a woman’s hand. It is therefore possible that our observed gender differences in vicarious touch were influenced by women in our study identifying more strongly with the observed touch compared to men. Future research should examine whether observer-touchee gender congruency influence the strength and quality of vicarious touch experiences.

### Emotional content & hedonic quality

The emotional content and hedonic quality of the observed touch strongly influenced the frequency and intensity of vicarious sensations. From prior research ^9–13,28^, we expected more intense sensory responses to videos depicting painful or threatening touch. Consistent with this, videos depicting painful interactions—characterised by high subjective arousal and perceived threat ^15^— elicited the strongest vicarious sensations. These were followed in intensity by reactions to pleasant and unpleasant touches, with neutral touches provoking the weakest responses. Our analysis revealed a strong correlation between the subjective arousal ratings from the VTD and the frequency and intensity of reported vicarious sensations, while the correlation with threat ratings was strong when considering only unpleasant and painful videos, due to the lack of range in threat ratings for pleasant and neutral videos. Our study relied on videos from the VTD, which includes assessments of subjective arousal, similar to other validated stimulus databases ^77–79^. We did not measure the physiological arousal induced by these stimuli (e.g., ^32^), but complementary physiological measures (e.g., skin conductance, heart rate) would be useful for future research.

When participants categorised their vicarious touch experience as neutral, pleasant, unpleasant, or painful, about half of the time, these responses matched the hedonic quality of the observed touch. However, two key deviations emerged. First, although painful videos elicited the highest intensity for vicarious sensations, participants often labelled their experience as unpleasant rather than painful. This may reflect a reluctance to use the most extreme response option, with ‘painful’ reserved for only the most intense experiences. Alternatively, it could indicate a genuine reduction in perceived vicarious pain intensity compared to the observed touch, as these videos were consistently categorised as showing painful touch by a separate sample of participants (VTD ^15^). Second, when participants’ vicarious touch reports did not align with the observed hedonic quality, the felt touch was most frequently categorised as neutral. This suggests that vicarious touch experiences may be lower in hedonic intensity overall. The absence of visible emotional reactions in our videos might have also reduced the empathetic engagement of some participants, potentially influencing their responses. Future research could explore the factors influencing how individuals classify vicarious sensations, particularly in relation to pain and general discomfort. Collectively, these findings highlight the role of emotional components in shaping sensory simulations of observed touch in the general population.

### Phenomenology & tingling

Our findings reveal distinct patterns of vicarious sensations across the four hedonic video categories, closely reflecting the type of touch depicted. Neutral videos, often featuring a single touch with moderate pressure, predominantly evoked ‘touch’ and ‘pressure’ sensations. In contrast, pleasant videos, which involved gentle strokes with soft objects or hands, most commonly elicited ‘ticklish’ and ‘tingling’ sensations. Thermal characteristics of touch were also reflected in participants’ experiences, with metal objects inducing ‘cold’ sensations and softer materials or another hand evoking ‘warm’ sensations. Consistent with previous research ^8,10,13^, ‘tingling’ was the most frequently reported sensation overall, occurring more often in response to pleasant videos than to painful ones, despite the latter’s higher threat and arousal ratings. This suggests that tingling sensations may mirror certain physical properties of observed touch, while likely also being influenced by cognitive factors such as attention, arousal, or heightened body awareness ^19,20,60,61^. Notably, tingling reports were more prevalent among individuals with fewer and weaker vicarious touch responses (see also ^10^), suggesting that diffuse, non-specific activation of the somatosensory mirror system may underlie this experience ^3^. The brain may interpret these weak or ambiguous sensory signals as tingling when they are insufficiently strong or coherent to be perceived as a distinct touch sensation^19^.

Vicarious sensations were most frequently reported in the hands, fitting with the location of observed touch. However, some participants reported sensations in other body regions, particularly in response to painful videos. These reports were concentrated in the upper body (e.g., arms, chest, back, stomach), suggesting a more spread-out sensory activation rather than strict somatotopic mirroring. One possible explanation is that emotionally salient touch engages interoceptive and autonomic networks involved in bodily awareness and threat detection, leading to a broader distribution of sensations. This aligns with findings that interoceptive awareness can influence vicarious touch experiences ^41,80^.

Another consideration is that the videos in our study depict self-touch (i.e., someone touching their own body) rather than interpersonal touch (see also ^81^). Neural evidence suggests that self- and other-touch may be processed differently ^39,82^, making it plausible that vicarious experiences of self-touch may also be distinct. Additionally, observers’ experiences can be shaped by whether they adopt the perspective of the toucher or the touchee ^41^. As our stimuli featured one hand touching another, we cannot distinguish whether participants identified more with the hand applying or receiving the touch. Future studies comparing the current VTD videos with matched videos featuring touch delivered by another person could determine whether these patterns extend across different types of observed touch.

Combined, our findings show that vicarious sensations often mirrored the observed touch, and most reported sensations were localised to the hands, suggesting genuine vicarious touch experiences. However, some reports may also reflect other affective or sensory processes. Our use of predefined response options aligns with prior research and enables systematic comparisons. While the ‘other’ option allowed flexibility, predefined choices may have influenced how participants interpreted and reported sensations, potentially shaping their responses or creating an expectation to report a sensation.

### Touch on objects

Observing touch on objects also evoked vicarious touch sensations, similar to those reported when observing touch on a hand, but at a much lower frequency. Holle et al. (2011) suggest this reduced response may stem from empathy toward the recipient of the touch, which would be absent in the object condition. Our findings align with prior research showing that individuals, including mirror-touch synaesthetes, sometimes report vicarious touch when observing touch on an object ^9,10,28,29,45^, and with EEG work demonstrating corresponding neural effects ^83^. This indicates that the mere observation of touch—whether on an object or a hand—can produce vicarious sensations.

### Different vicarious touch profiles

Using a data-driven clustering approach we tested whether there exist distinct variations in vicarious touch perception within the general population, similar to what has been previously done for vicarious pain ^12^. We clustered vicarious touch responders based on the total endorsements across the 40 video trials (1-40), the total number of trials with a reported exteroceptive sensation (0-40), and the total number of trials with a reported interoceptive sensation (0-40). Our results indicated meaningful clustering into three distinct groups, suggesting that there exists different sensory response profiles in the general population, rather than vicarious touch being a uniform phenomenon, which is similar to findings on vicarious pain ^12^.

The distribution of exteroceptive and interoceptive sensations differed across groups, with those who frequently reported vicarious touch endorsing relatively more exteroceptive than interoceptive sensations. This pattern is consistent with previous research suggesting a link between stronger vicarious responses and more externally localised sensations ^10^. Plausibly, individuals with stronger vicarious responses may more precisely simulate the tactile qualities and spatial location of the observed touch, while those with weaker responses may experience more diffuse or internally directed sensations ^3,8,10,19^. However, while we classified sensations based on prior work ^10^, the distinction is not always clear-cut ^84^. Sensations such as tingling and tickling may sometimes be perceived as externally generated ^19^, blurring the boundary between exteroception and interoception. Future research should investigate whether these categories correspond to distinct neural mechanisms or instead reflect a continuum of sensory experiences shaped by individual differences in somatosensory processing.

In addition to differences in total vicarious touch endorsements and interoceptive and exteroceptive sensations used for clustering, the clusters predicted other differences. Individuals in Cluster 3 (strong responders) reported hand-localised sensations more frequently than those in Cluster 2 (medium responders) and Cluster 1 (weak responders), suggesting a more specific mirroring in this group. Compared to Cluster 1, members of Clusters 2 and 3 exhibited higher intensity across all sensation types, indicating that a higher tendency for vicarious touch correlates with increased sensation intensity. However, the overall intensity reported by Cluster 3 was not greater than that of Cluster 2, suggesting that vicarious touch intensity may plateau at moderate levels of responding. The exception was the intensity of reported pain, which was highest in Cluster 3. This implies that the intensity of mirroring painful touch might have a higher threshold before reaching a plateau. While earlier studies found greater differentiation in individuals with higher vicarious touch endorsements when observing touch to another person versus an object, measured in intensity ^10,28^, our findings revealed no such distinction across clusters. Interestingly, emotional reactivity scores were also consistent across all clusters, indicating that the tendency to experience vicarious touch might be distinct from general emotional empathy (see also ^4,45^), which would contradict some previous findings ^8,10,29^.

Most knowledge about vicarious touch comes from research on mirror-touch synaesthetes ^5,8–10,28–33^. Some individuals in our sample may exhibit characteristics of mirror-touch synaesthesia. Those in Cluster 3 frequently reported intense, localised, exteroceptive sensations rather than generalised, interoceptive sensations. However, these individuals did not show a clearer distinction in their responses to human versus object touch, nor did they exhibit enhanced emotional reactivity, previously associated with mirror-touch synaesthesia ^10,29,46^. Cluster 3 comprised 10.4% of our entire sample (12% of all vicarious touch responders), closely matching the prevalence of mirror-touch synaesthesia when relying on self-report measures ^8,85^. However, prior behavioural work using a visual-tactile interference task found a much lower prevalence of 1.6% ^8^. When we looked at an individual level, there were six participants in Cluster 3 (1.4% of our entire sample and 1.7% of vicarious touch responders) who consistently reported exteroceptive sensations (>36/40 trials) localised to the hands (100% of endorsed trials), effectively appearing as outliers in the sample. It may be the case that these six individuals experience mirror touch synaesthesia. Future research could combine our approach with objective measures, such as a visual-tactile congruency task ^8,29^ or neuroimaging (e.g., see ^4^) to determine whether individuals in a specific cluster, including any outliers, exhibit behavioural or neural differences consistent with mirror-touch synaesthesia.

Our findings collectively indicate that consciously experienced vicarious touch is not a rare phenomenon, at least in experimental settings, and extends beyond conditions such as mirror-touch synaesthesia ^8,10,34,86–88^. The high prevalence observed in our study supports the idea that vicarious touch exists on a continuum, with mirror-touch synaesthesia representing one extreme ^7,34–36^. However, our results also suggest qualitative differences in vicarious touch experiences across individuals, akin to the distinct response profiles reported for vicarious pain ^12^. These variations suggest that individual differences in certain aspects, such as interoceptive sensitivity, sensory processing, or attentional focus, may shape how vicarious touch is experienced ^19,20,60,80,89,90^. Recognising these diverse response patterns may offer deeper insight into the mechanisms underlying sensory mirroring in the general population.

Our broad investigation lays the groundwork for future research, which could extend to a broader and more diverse population sample, including various age groups and cultural backgrounds. Additionally, it would be valuable to study the sensory and emotional characteristics of vicarious touch in populations with unique sensory experiences, such as individuals on the autism spectrum ^42,91,92^, those experiencing depersonalisation ^93^, or individuals with ASMR ^90^. Testing these diverse groups could provide deeper insights into the variability of vicarious touch, potentially informing therapeutic strategies for those with atypical sensory responses ^40,42,92,94^. Profiling individuals based on their vicarious sensory perception may also yield valuable information on how best to support professions where practitioners often deal with others’ pain, such as healthcare professionals, thereby optimising care while protecting the well-being of caregivers ^95,96^.

## Conclusion

Here, we examined vicarious touch reports in a general student sample using stimuli from the Validated Touch-Video Database (VTD ^15^). We found that a large majority of participants reported experiencing localised sensations while observing touch on a hand, with these sensations occurring more frequently in women than in men. Notably, there was no correlation between vicarious touch responding and emotional reactivity scores, suggesting that the capacity for sensory mirroring may be distinct from emotional disposition. The type of sensation reported—such as tingling, pressure, or pain—often matched the physical characteristics of the observed touch, reinforcing the link between visual input and sensory response. Additionally, the emotional content of video stimuli, such as the hedonic quality and level of subjective arousal, played a key role in shaping vicarious experiences, with painful touch eliciting the strongest sensations. Using a data-driven clustering approach, we identified distinct vicarious touch response profiles. Individuals who reported more frequent vicarious touch experiences tended to describe tactile, localised sensations in the hands that directly mirrored the observed touch, whereas those with fewer vicarious touch responses were more likely to report diffuse or generalised sensations. Our findings highlight the widespread nature of vicarious touch, supporting the idea that these experiences exist on a continuum, with mirror-touch synaesthesia at one extreme, while also revealing individual differences in how they are experienced. By identifying how the sensory and emotional aspects of observed touch influence vicarious touch experiences, this research contributes to a broader understanding of how we perceive and simulate the tactile experiences of others.

## Data availability

All video stimuli, analysis scripts and data can be found online (https://osf.io/npqu3/).

## CRediT authorship contribution statement

**Sophie Smit:** Conceptualisation, Methodology, Project administration, Data curation, Formal analysis, Visualisation, Writing – original draft, Writing – review & editing. **Matthew J. Crossley:** Conceptualisation, Methodology, Formal analysis, Visualisation, Writing – review & editing. **Regine Zopf:** Conceptualisation, Writing – review & editing, Supervision. **Anina N. Rich:** Conceptualisation, Methodology, Writing – review & editing, Supervision.

## Additional information

This work was supported by a Commonwealth funded Research Training Program and Macquarie University Research Excellence Scholarship awarded to SS. ANR is supported by an Australian Research Council Future Fellowship (FT230100119). Authors declare no competing interests.

