## Supplementary materials for "Characteristics of vicarious touch reports in a general population"

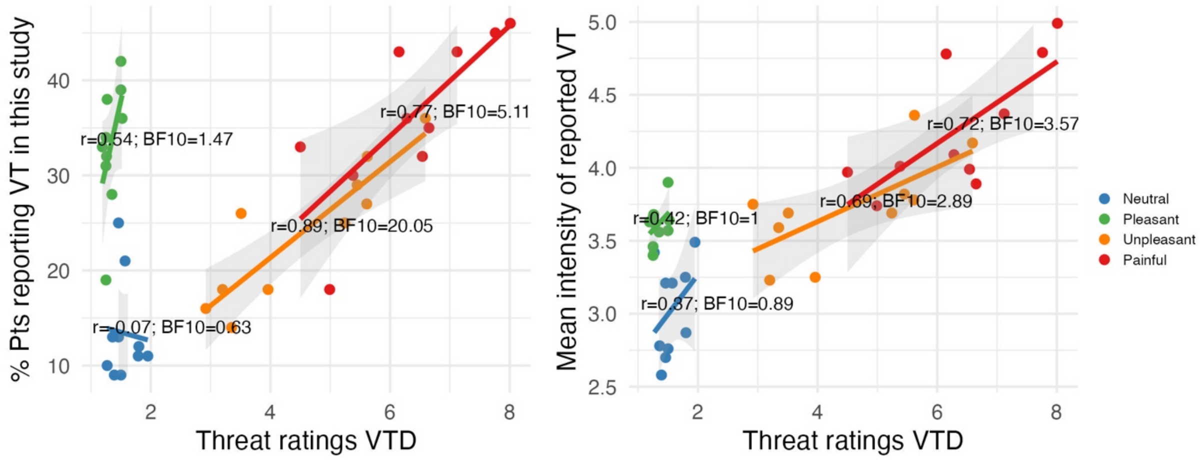


**Supplementary Figure 1. Correlation between the threat ratings from the VTD and percentage of participants that endorsed vicarious touch (left) and mean intensity (right) for each hedonic category.** Scatterplots depict the relationship between the independently rated threat on a scale from 1-10 (1 = not at all, 10 = extremely) from the VTD and the endorsement (left) and intensity (right) of evoked vicarious touch for each video. Each dot signifies the percentage of participants that endorsed vicarious touch, or mean intensity ratings across participants, for the individual videos, color-coded by category: neutral, pleasant, unpleasant, and painful, with 40 videos in total. Individual lines show linear regression for each category with shaded 95% confidence intervals. VTD = Validated Touch-Video Database (Smit & Rich, 2023); VT = Vicarious touch. BF = Bayes factors.

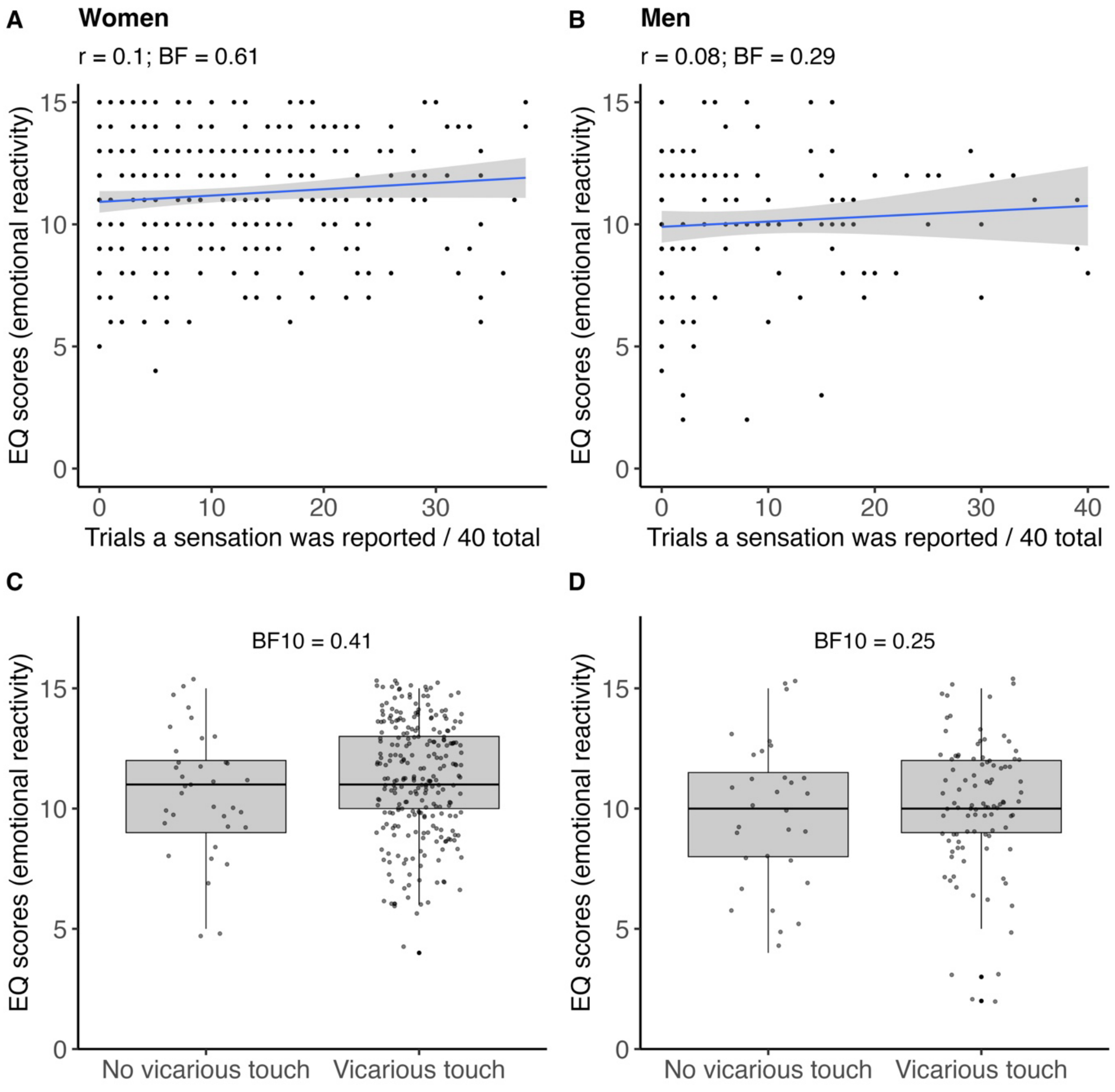


**Supplementary Figure 2. Vicarious touch endorsements and emotional reactivity scores by gender.** Separate plots are shown for women (left) and men (right). (Top) Scatterplots depict the relationship between the number of vicarious touch endorsements out of 40 trials total and emotional reactivity scores based on five questions from the EQ (higher scores suggest higher emotional empathy). Lines show linear regression with shaded 95% confidence intervals. (Bottom) Comparison of emotional reactivity between participants with no vicarious touch and at least one vicarious touch response. Scatterplots show the distribution of scores for the two groups. The line within each box indicates the median, the box boundaries indicate the interquartile range, and the whiskers extend to 1.5 times the interquartile range.

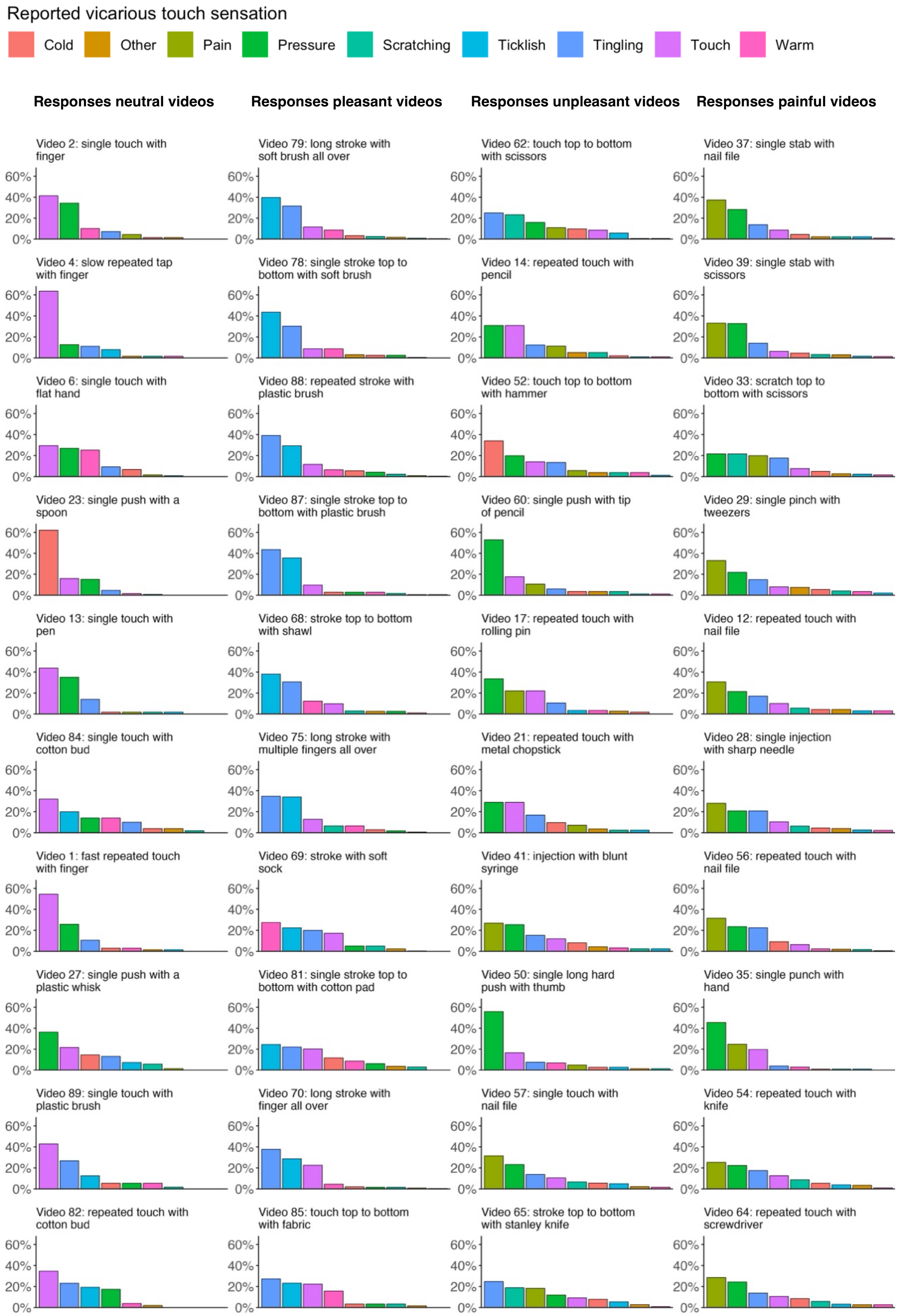


**Supplementary Figure 3. Detailed sensation distributions across hand-touch videos, highlighting the influence of touch method and type of object or hand.** The relative frequencies of different vicarious touch sensations reported by participants in response to watching 40 videos selected from the VTD (note video numbers relate to the code given in the VTD). Each column represents a different category of video: neutral, pleasant, unpleasant, and painful. For each video, sensations are sorted in decreasing order of their reported frequency. Sensations are represented using different colours, and their definitions can be found in the accompanying legend. The y-axis displays the percentage of each sensation type relative to all reported sensations across the ten videos within each of the four categories.

Considering the sensations reported for each video provides insights into nuanced touch interactions and their corresponding sensations. For example, in two videos, touch was applied by scissors moving from the top to the bottom of the hand. When touch was applied with more pressure (video 62; pain category), most participants reported feeling pressure, whereas when touch was applied softly (video 33; unpleasant category), the most common response was tingling. In both cases, the second most common response was scratching, reflecting the scratching across the hand. In addition, we found that pleasant videos evoked the highest percentage of warm sensations whereas unpleasant videos evoked the highest percentage of cold sensations. This appears to capture anticipated sensations from the object that is applying the touch in the videos. For example, most of the unpleasant videos showed a ‘cool’ object applying the touch, including metal items such as a hammer, spoon, knife, scissors, or screwdriver, and this often resulted in cold sensations (video 23; neutral category, video 52; unpleasant category). Touching objects in the pleasant videos included soft or warm items such as a soft brush, cotton pad, a piece of fabric and a flat hand, and this commonly resulted in corresponding warm sensations (videos 69 and 85; both pleasant category). This shows that variation in pressure and the type of object used in the visual touch stimuli influences the specific character of the vicarious touch sensation.

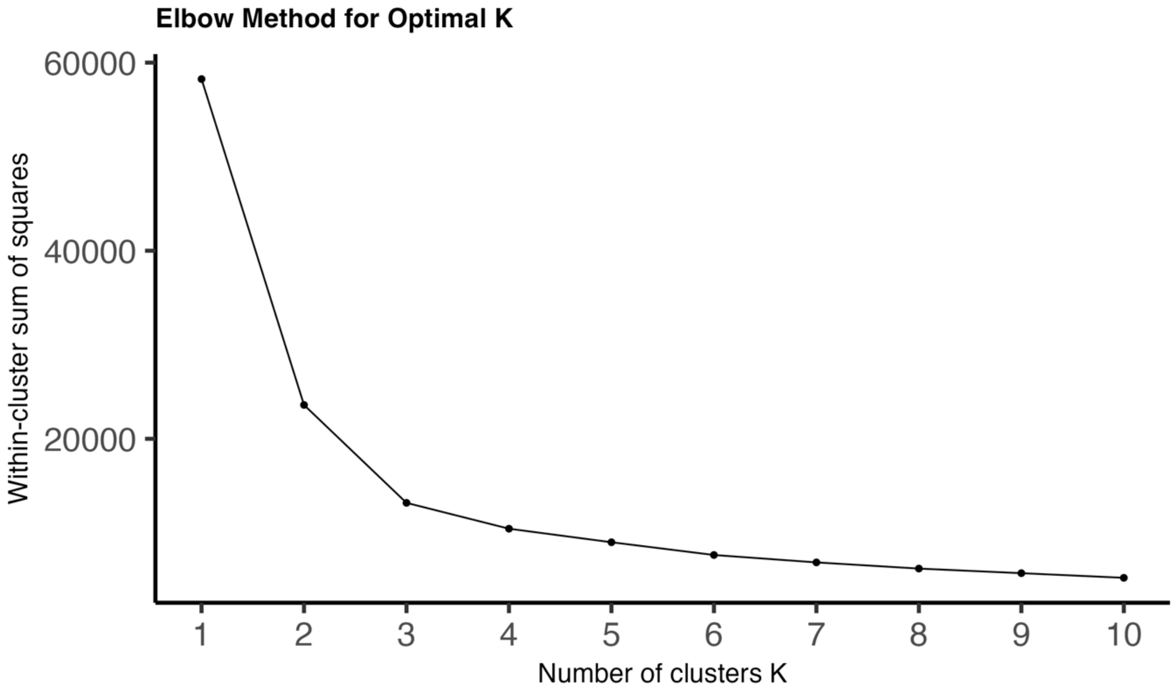


**Supplementary Figure 4. Elbow method for determining optimal number of clusters.** The plot illustrates the within-cluster sum of squares (WCSS) against the number of clusters (K), ranging from 1 to 10. Each point represents the total variance within the clusters for a given K, with a line connecting the points to highlight the trend. The "elbow" in the plot, where the rate of decrease in WCSS shifts notably, suggests the most appropriate number of clusters for k-means clustering. This method helps to identify a suitable K by selecting the point at which increasing the number of clusters results in diminishing returns on explained variance (James et al., 2013; MacQueen, 1967).

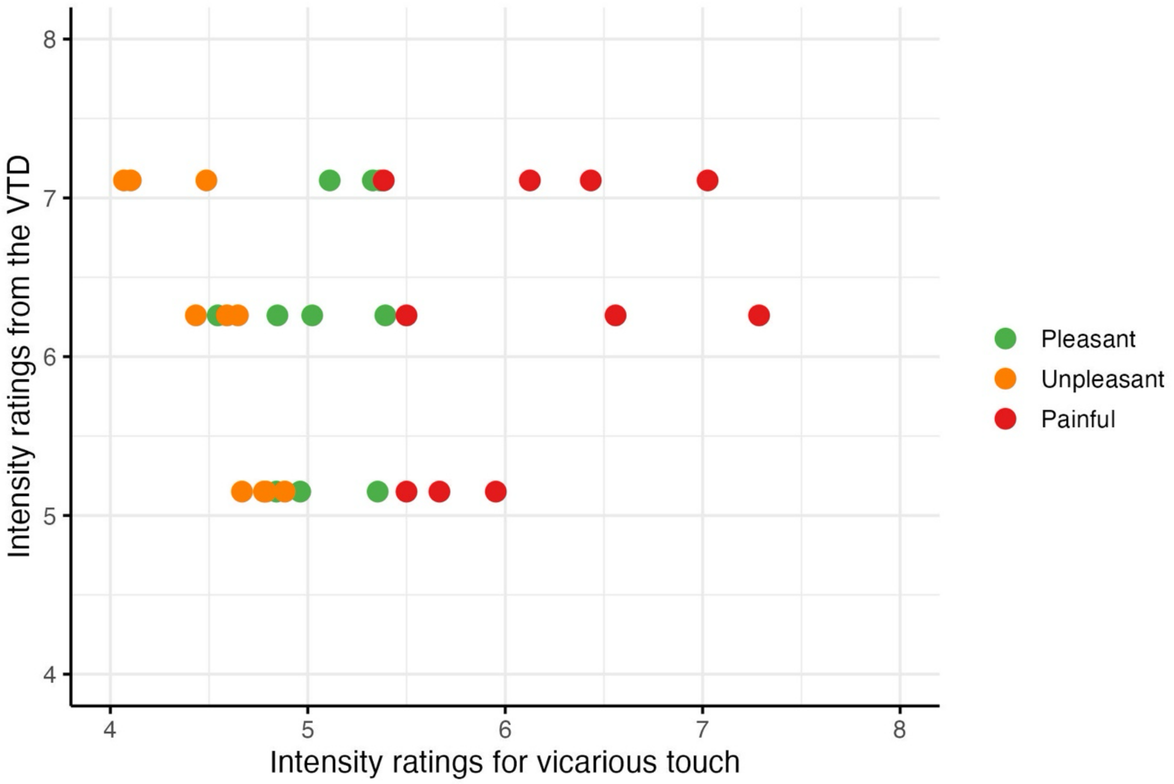


**Supplementary Figure 5.** The relationship between the hedonic intensity ratings for vicarious touch in the current study (i.e., for felt touch) and the hedonic intensity ratings from the VTD (i.e., for seen touch) across three categories: pleasant, unpleasant, and painful. Each point represents the average intensity rating for a specific video, with different colours indicating different hedonic categories. The plot illustrates how participants' felt hedonic intensity of vicarious touch correlates with the observed hedonic intensity of touch in the videos.
